# Unleashing alternative polyadenylation analyses with REPAC

**DOI:** 10.1101/2022.03.14.484280

**Authors:** Eddie L Imada, Christopher Wilks, Ben Langmead, Luigi Marchionni

## Abstract

Alternative polyadenylation (APA) is an important post-transcriptional mechanism that has major implications in biological processes and diseases. Although specialized sequencing methods for polyadenylation exist, their presence in public repositories is extremely limited when compared to traditional RNA-sequencing. To overcome this, we developed REPAC, a framework for the analysis of APA from RNA-sequencing data. REPAC implements a new method for detection of APA and is designed to take advantage of *recount3* which enables a streamlined way to analyze over 750,000 publicly available samples. Using REPAC, we investigated the landscape of APA caused by activation of B cells. Our analysis revealed that during this process, hundreds of genes are regulated by APA, most notably genes involved in the secretion pathway which is central for the transition to antibody-secreting B-cells. Moreover, we also showed that many genes associated with interferon response are also shortened, suggesting that APA might also play a significant role in the immune response. We also show that REPAC is faster than alternative methods by at least 7-fold and that it scales well to analysis involving hundreds of samples. Overall, the REPAC method offers an accurate, easy, and convenient solution for the exploration of APA across many phenotypes.

## Background

Mechanisms that control gene expression at the RNA level are often referred to as post-transcriptional regulation (PTR) mechanisms. Splicing and alternative polyadenylation (APA) are well-known examples of PTR that can regulate not only gene expression but also their function. While splicing has been extensively studied since the advent of Next Generation Sequencing (NGS), APA studies are far less common than splicing studies. Indeed, inferring APA events from RNA-Seq data is challenging due to the lack of an intrinsic characteristic (e.g., split-reads for splicing) and for this reason, several specialized sequencing methods were developed to pinpoint polyadenylation sites (PAS) [1–3]. Although these methods improve the quantification of PA sites usage, the number of publicly available data derived from these methods is extremely limited in comparison to traditional RNA-Seq data.

This poses a challenge to the study of polyadenylation (PA) biology given that the lack of publicly available data severely hinders investigation and hypothesis generation without incurring major experimental costs. To overcome this limitation, several groups have developed methods to quantify PA usage from RNA-Seq data [4–6]. Among them, DaPars [6] is a popular method to compare PA profiles acrosstwo phenotypes and detect the differential usage of APA by the degree of difference in APA usage quantified as a change in Percentage of Distal polyA site Usage Index (*ΔPDUI*). Likewise, QAPA [4], which leverages the speed and power of pseudoalignment software, such as salmon, was shown to be the fastest method for APA studies so far.

Most of the methods currently available rely on the effect size of the change in the proportion of the expression levels between different PA sites. Performing statistical analysis on proportions of a total (i.e. percentage) imposes many statistical limitations that are often ignored and can lead to inaccurate results. Most multivariate methods that were developed for real values cannot be directly applied to compositional data (proportional data) because compositional data often breaks many assumptions of these methods [7]. Moreover, these methods do not allow for the control of unwanted variables or the design of more complex comparisons (e.g., factorial designs, paired samples, etc.).

In this work, we present Regression of PolyAdenylation Compositions (REPAC) a novel framework to detect differential alternative polyadenylation (APA) using regression of polyadenylation compositions which can appropriately handle the compositional nature of this type of data while allowing for complex designs. We show that REPAC is faster and yields more accurate and robust results in comparison to other methods.

## Results

### REPAC can accurately detect APA events

REPAC makes use of expression estimates of 50bp windows upstream of annotated PA sites to fit a generalized linear regression model on the compositions to assess differential polyadenylation site usage (DPU) between conditions. This quantification can be done with traditional tools (e.g., Subread, HT-Seq, etc.) or directly pulled from *recounts*[8] bigWig files (Figure 1). Briefly, to assess the performance of REPAC, we simulated 2 conditions (n=5 for each condition) with 5000 genes having a longer or shorter isoform. Half of the set had one of the isoforms predominantly expressed in one of the conditions (1250:1250 for short/longer isoforms) and the other half did not have a predominant isoform. The ratio of longer to shorter isoforms was randomly defined within a pre-defined range (see methods). We applied the REPAC method after removing 197 low expressed genes. For each type of event, REPAC was able to achieve 0.99/0.98, 0.99/0.98, and 0.98/0.98 specificity/sensitivity for lengthening, shortening, no preference (NP), respectively. With an overall accuracy of 0.98 and an area under the curve (AUC) of 0.9989, REPAC was shown to accurately detect APA events (Figure 2A).

**Figure 1.**
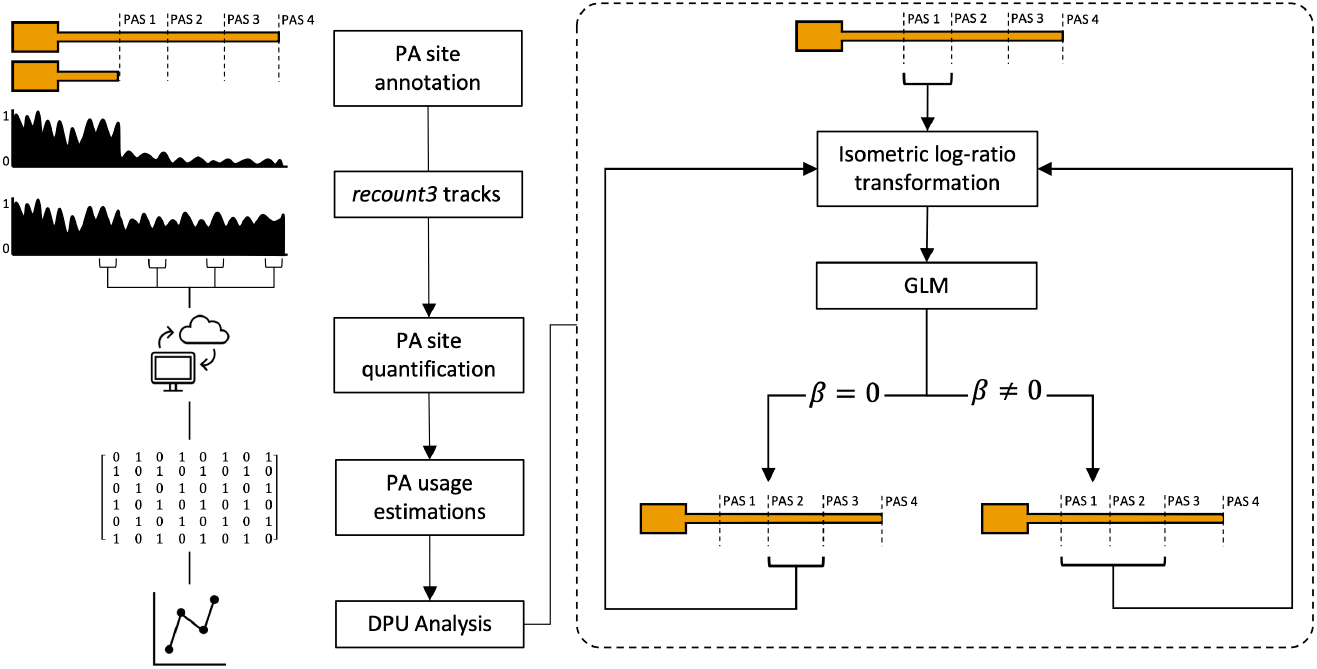
REPAC workflow. REPAC perform analysis of differential polyadenylation usage by analyzing the upstream region of annotated PAS. While quantification of PAS can be performed in traditional ways (alignment and counting), it was primarily design to take advantage of the *recount3* project to extract counts on-the-fly for over 750,000 samples publicly available.

**Figure 2.**
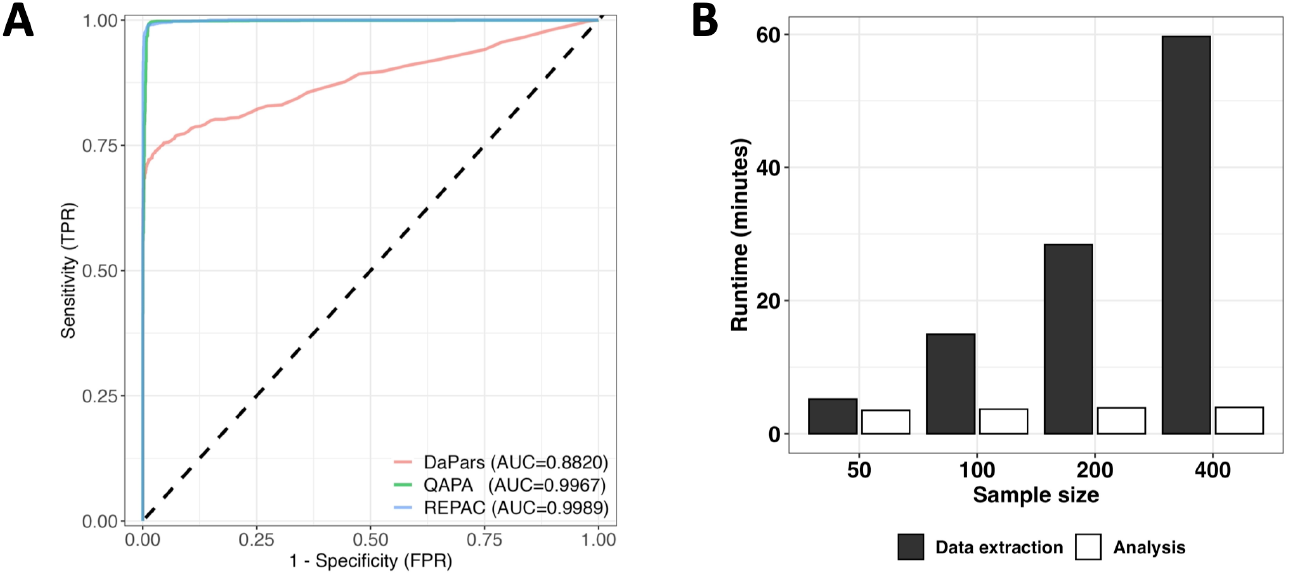
Benchmarked tests. A) Empirical ROC curves of REPAC, QAPA and DaPars. REPAC and QAPA performs very similarly (*AUC* = 0.99) in simulated data, while DaPars shows a lower performance (AUC = 0.88). Curves were based on the effect-size of the methods i.e., cFC, *ΔPDUI* and ΔPAU for REPAC, DaPars and QAPA, respectively. B) REPAC runtime scales almost linearly with the number of samples.

### Performance of alternative methods for APA detection

We compared the performance of REPAC to other popular methods for APA detection: DaPars[6] and QAPA[4]. QAPA relies on external tools such as salmon to obtain estimates of each isoform 3’-UTR expression level which are transformed to relative expression levels by normalizing the isoform expression level by the total 3’-UTR expression level. In contrast, DaPars relies on 3’-UTR changes in coverage to infer *de novo* PA sites and their differential usage.

We measured the performance of all the methods using the same simulation experiment. For each method, we used their respective effect size estimators (*ΔPDUI* for DaPars and *ΔPPAU* for QAPA) to compute the AUC similarly to what done for REPAC. Out of the three methods tested, REPAC and QAPA were able to achieve high accuracy, with REPAC presenting a marginally higher performance (AUC = 0.9989 *versus* 0.9967), while DaPars exhibited the lowest performance in the simulated dataset (AUC = 0.882) (Figure 2A). The lower performance of DaPars was on par with previous simulations suggesting that *de novo* prediction of PAS from traditional RNA-seq data is relatively inaccurate compared to methods that rely on annotated PAS [4]. Moreover, DaPars was drastically slower (> 40-fold) than REPAC and QAPA.

### REPAC enables fast and streamlined exploration of public domain data

The REPAC method and associated R package were designed to take advantage of our recently published recount3 resource[8]. The recount3 project processed and analyzed data from over 750,000 samples of human and mouse origin publicly available in SRA in a standardized manner. The REPAC package extends recount3 to enable the extraction of expression estimates for thousands of PA sites within a few seconds per sample. The REPAC framework has major advantages over existing methods for APA analysis from RNA-seq data. For instance, it allows the user to skip the time- and storage-intensive step of downloading the raw data and processing it. By integrating with the recount3 framework, REPAC enables researchers to explore differential APA events for thousands of phenotypes in a fast and streamlined fashion. QAPA has already been shown to be substantially faster than other methods such as DaPars, ROAR, etc. Therefore we benchmarked the speed of REPAC against QAPA. To this end, we started by obtaining from SRA the raw data for the project SRP048707 (see below). The total size of the data set analyzed was approximately 27 GB (gzipped fastq files). Assuming a constant speed of 300Mbps, this step alone would take approximately 15 minutes to complete. Because REPAC pulls expression estimates directly from recount3 it allows the users to skip this step. Next, pre-processing (extracting 3’-UTR sequences, building the index and quantification) and expression quantification with salmon[9] for QAPA analysis took on average 17 minutes, while obtaining PAS estimates with the REPAC package took under one minute. Finally, detecting DPU between the comparisons took 0.03 minutes with QAPA versus 3.2 minutes with REPAC. However, it is important to note QAPA does not perform any statistical testing, which contributes to the speed in this step. Overall, considering the entire workflow of an APA study, REPAC was 7.6 times faster than QAPA.

### The REPAC method is scalable to large data sets

One advantage of REPAC is its ability to tap into recount3 to directly obtain PAS expression quantification. As demonstrated in the previous analysis, this feature can greatly speed up the analytical process even for small data sets. However, the advantages of REPAC become abundantly clear when analyzing large collections, such as the GTEx for instance, for which storage and computing power requirements can quickly become a limiting factor for many researchers. Moreover, the process of acquiring access to raw data can be slow and burdensome. To test how well REPAC scales with increasing amounts of data, we extracted PAS quantification and performed a DPU analysis for a total of 20, 100, 200, and 400 randomly selected brain (cortex) and testis tissue samples from the GTEx project (V8). We found that the time to quantify PAS with REPAC scaled linearly with the number of samples, with an average of 9 seconds per sample. The time to test for DPU between conditions remained stable at around four minutes (Figure 1B).

When looking at highly differential PA site usage (*|cFC*| ≥ 0.25, *FDR* ≤ 0.05) between 200 testis and 200 brain samples, we observed 879 genes with differential usage of PAS. The results showed that testis favors the expression of shorter 3’-UTR isoforms when compared to the cerebral cortex, with over 97% of the genes detected showing preferential usage of a shorter PAS in testis and vice-versa. These results were consistent with previous studies reports [10, 11].

### Global profiling of APA during B cell activation

We applied the REPAC method to investigate the landscape of APA in response to B cell activation. To this end, we used a dataset from Diaz-Muñoz et al[12] containing naive and LPS-activated B cells (with four replicates per condition). Using the REPAC package, we obtained expression estimates for 67,509 3’-UTR PAS (as derived from the PolyASite 2.0 database[13], see methods) for all samples, and then performed a DPU analysis comparing naive *versus* LPS-activated B cells. This analysis detected 117 genes with DPU in response to B cell activation (*|cFC*| ≥ 0.25, adjusted p-value ≤ 0.05). Approximately 80% of the genes with DPU were found to have a higher usage of a more proximal PAS upon B cell activation, with the 3’-UTR being 817 bp shorter on the median. We found that genes associated with secretion mechanisms, such as Cd47, Edem1, and Rbx1, were among the most significant DPU events (Figure 3A). Next, we investigated whether these changes were associated with a particular biological process (BP). Through gene set enrichment analysis (GSEA) of GO BP, we observed that 19 BP were significantly enriched in 3’-UTR-Shortening (3’-US) events (adjusted p-value≤ 0.05). Among the top enriched pathways were IRE1 mediated unfolded protein response and response to type I interferon (IFN-I) (Figure 3B). Some of these findings were consistent with findings from Cheng and collaborators[14] showing that in activated B cells, the genes associated with secretion exhibit shorter 3’-UTRs. Notably, our analysis also expanded the currently known set of genes and processes affected by APA - revealing processes, such as response to IFN-I, that have not been associated with B cell activation before.

**Figure 3.**
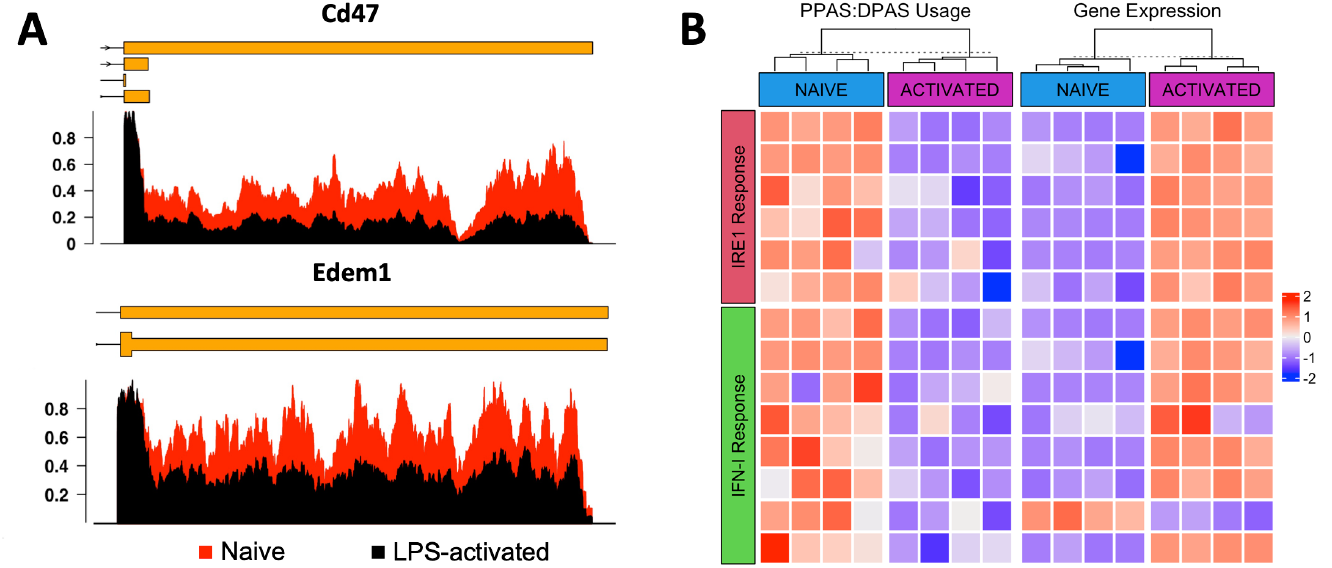
3’-UTR-shortening events in naive and LPS-activated B cells. A) Differences in 3’-UTR coverage of secretion associated genes Cd47 and Edem1. B) Heatmap showing the relationship between the usage of the proximal and distal PAS (PPAS:DPAS) and expression levels (CPM) using stardardized values. Overall, preferential usage of the shorter 3’-UTR isoform (blue color) is associated with higher expression levels (red color) for the genes involved in secretion and IFN-I response.

Additionally, we also performed the same comparison using QAPA to evaluate the performance of both software in real data. The analysis by QAPA was able to detect 13 genes with DPU (*Δ|PPAU*| ≥ 20), of which approximately 53% were 3’-US events. GSEA analysis on QAPA results did not indicate enrichment in any of the processes found enriched by REPAC, including secretion pathways previously reported by other studies[14]. Surprisingly, none of the events detected by REPAC were captured by QAPA and vice-versa. Given this discrepancy, we visually evaluated inspect the results for the top 10 predicted events. Upon visual inspection of the results, we found that the events predicted by QAPA were largely driven by false positives caused by low 3’-UTR coverage, and no striking changes were observed in genes with enough coverage (Additional File 1), while the events predicted by REPAC clearly showed a difference in 3’-UTR coverage (Additional File 2).

### Technical implications of APA in downstream analyses

Our results indicated that APA events in 3’-UTR regions can drastically impact gene size. Therefore, we investigated how APA events can impact downstream steps, such as differential gene expression analysis. To this end, we obtained length-corrected (length scaled TPM) and raw gene expression estimates via the salmon-tximport pipeline and carried out a differential gene expression analysis between naive and activated B cells. After taking into account the different lengths of the isoforms, we found that many genes with APA events captured by REPAC exhibited significant changes in expression levels (Figure 4A) and fold-change between conditions (Figure 4B).

**Figure 4.**
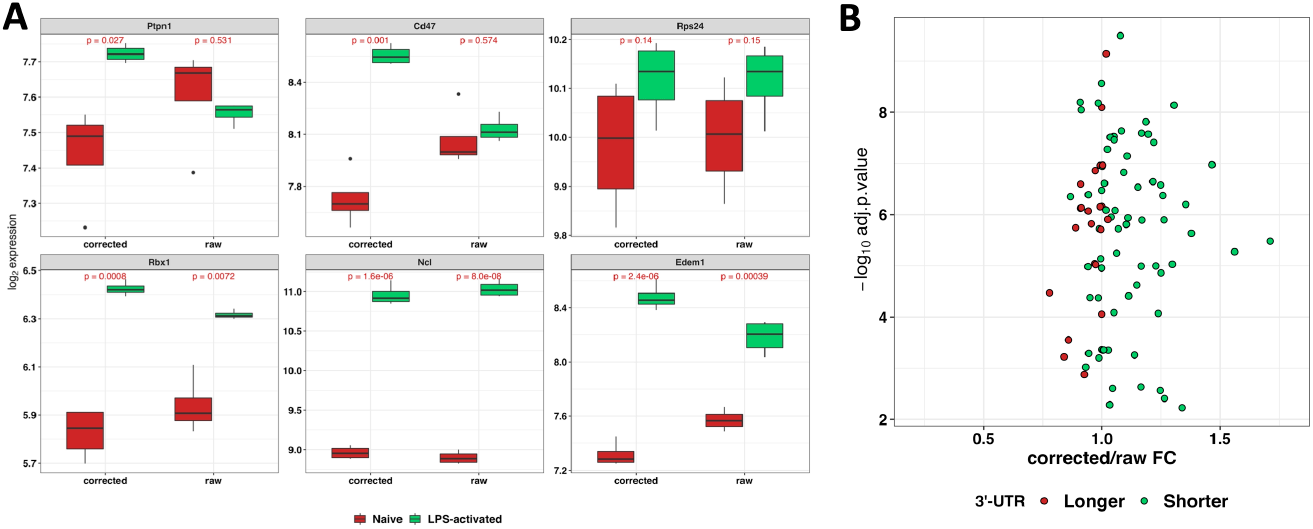
APA can drastically impact expression estimates. A) Expression levels with and without length correction by tximport for the top 6 APA events predicted by REPAC. Many genes can be found deferentially expressed after length correction, highlighting the importance of APA in other analysis. P-values from a t-test between groups are shown. B) Scatter plot shows the ratio between fold-change estimated from a differential gene expression analysis with and without length correction.

This issue was evidenced by genes such as Cd47, Ptpn1, and Edem1, whose expression levels after LPS-activation increase by 75%, 222%, and 20%, respectively, when adjusting for the transcript length. In contrast, if the transcript length is not taken into account, the observed changes in expression level upon LPS-activation for Cd47, Ptpn1, and Edem1 are 2%, 151%, and −4.7%, respectively (Figure 4A).

Genes predicted by REPAC to have 3’-US upon B cell activation showed predominantly increased fold-change values, while genes predicted to have 3’-UTR lengthening (3’UL) presented a decrease in fold-change when comparing fold-changes before and after isoform length correction (Figure 4B). For some genes, we did not observe significant changes in fold-change after correction due to alternative PAS for these genes not being annotated as independent isoforms meaning these differences are usually not captured by gene expression quantification software.

## Discussion

Here we present Regression of PolyAdenylation Compositions (REPAC), a new framework for the study of differential PA events using traditional bulk RNA-sequencing data. The REPAC method is based on the principles of compositional data (CD) analysis developed by Aitchson[7].

In our simulated data set REPAC outperformed DaPars, which is currently, one of the most popular methods of APA analysis, in both accuracy and speed. When compared to more recent developments such as QAPA, REPAC presented a marginal increase in performance (Figure 2A). However, when comparing the results of REPAC and QAPA in real RNA-seq data, we found that REPAC results were more accurate and robust than QAPA (Additional Files 1 and 2). Moreover, REPAC offers substantial advantages over other existing methods.

The REPAC package takes full advantage of the recount3 framework to skip the data acquisition and processing steps and directly extract the necessary data for differential APA analysis with little resource usage and time. This feature makes over 750,000 samples readily available for analysis (Figure 1). This is a huge advantage over existing methods since all of them require raw data to be obtained and processed before analysis. Even when disregarding the time and effort of data acquisition, DaPars and QAPA still require pre-processing of the data such as aligning the data to the genome or performing 3’-UTR estimations with salmon, respectively. Despite salmon being relatively lightweight and faster than alignment-based methods (i.e. DaPars), it still requires substantial computational resources to carry the index construction and quantification, especially when a large number of PAS are used. Additionally, since REPAC makes use of generalized linear regression models, it can easily handle complex comparisons (i.e. factorial designs, paired samples, etc.) and correct for unwanted sources of variations (i.e. batch effects, blocks, etc.), which cannot be directly modeled by other methods.

All of these overheads translate into a much faster and more accurate analysis with REPAC than other methods. We demonstrated that the analysis of a small data set (n=8) was 7.6 times faster to process with REPAC than QAPA (4.2 *versus* 32 minutes), which already outperforms other methods[4]. Even when disregarding the time for data acquisition, REPAC was still 4 times faster than QAPA. It is important to note that these differences in processing times would significantly scale with larger data sets, to the point where performing APA analysis with other methods, might become infeasible for research groups without access to a robust computing environment.

As a proof of principle, we used REPAC to compare brain (cortex) and testis tissues from GTEx V8. On average, REPAC took 9 seconds per sample to quantify all PAS, meaning REPAC is scalable for large data sets with 400 samples taking under one hour to complete the entire analysis (Figure 2B). In contrast, other methods would require the user to first get permission to access the raw data, download the data, and processes it, all of which would take days and access to a robust computing environment that are compliant with data privacy laws (i.e. Health Insurance Portability and Accountability Act in the US, and General Data Protection Regulation in Europe).

Moreover, the results of this comparison were consistent with previous studies reporting that cells from non-proliferative tissue (eg, brain) tend to express longer 3’-UTR isoforms than cells from proliferative tissues (eg, testis)[10, 11]. Interestingly, we found that 3’-US events were enriched for genes involved in spermatogenesis and androgen response, suggesting that APA is not only associated with a proliferative state but also regulates specific processes associated with tissue function.

Despite one of the first pieces of evidence of APA as a functional mechanism being reported during B cell transition to plasma cell [15], the landscape of APA during B cell activation had remained largely under-explored. Recently, a study by Cheng and collaborators [14] conducted a broad survey of APA in secretory cell differentiation and observed that many genes involved in secretion presented 3’-UTR shortening after B cell activation. Our analysis of naive *versus* LPS-activated B cells detected hundreds of genes impacted by APA, whose majority were 3’-UTR shortening events. On par with previous observations[14], we found that genes involved in secretion were enriched in 3’-UTR shortening. Specifically, we found that genes involved in the IRE1 mediated unfolded protein response enriched in shortening events (Figure 3B).

Interestingly, response to type I interferon (IFN-I) was among the most enriched processes in 3’-US. Early response of B cells to IFN-I has been shown to elicit many types of responses (e.g., enhance antiviral humoral feedback by increasing the formation of early antiviral IgM, increase TLR-9 mediated activation and regulate autoreactive B cell activation[16–18]). Using REPAC we were able to detect 3’-US in genes such as Ptpn1, Ube2k, and Irf2bp2, which are known to be involved in cell response to interferon stimulation[19–23]. To the best of our knowledge, this is the first report of regulation of IFN-I response in B cell by APA.

Finally, we also demonstrated that APA has serious implications on downstream analysis (e.g., differential gene expression analysis) since traditionally gene length is assumed to be the same between conditions and therefore is not accounted for in many approaches. While recent pipelines based on expression estimates at transcript level through pseudo-alignments, such as the salmon-tximport[9, 24] pipeline used in this analysis, can correct for differential isoform/3’-UTR usage, they are only able to do so if these isoforms/3’-UTRs are properly annotated. In this regard, we found that major annotation resources like GENCODE/ENSEMBL and RefSeq still lack proper annotation of 3’-UTRs for many genes. Therefore, the accessibility and ease of usage of REPAC is a powerful tool in making sure APA events are properly detected, annotated, and studied.

## Conclusions

We demonstrated that REPAC is a robust and powerful tool for exploring the biology of APA. It enables the analysis of over 750,000 samples encompassing thousands of different phenotypes. With REPAC, we want to encourage more studies on APA and how they influence normal and disease tissues. We hope this new tool can help pave the way to develop new hypotheses that can be further explored to understand the biological role of APA as a whole.

## Methods

### Detection of differential polyadenylation site usage

The REPAC framework is based on the Aitchison geometry in the simplex. A D-part simplex is defined as:

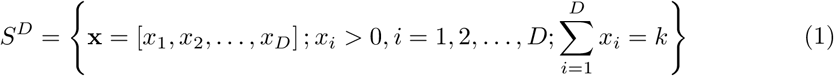

where D is the number of elements of comprising a composition (i.e., number of PAS) and *k* is a positive constant. We apply the isometric log ratio (*ilr*) transformation to the simplex. Let x be a D-part simplex, then:

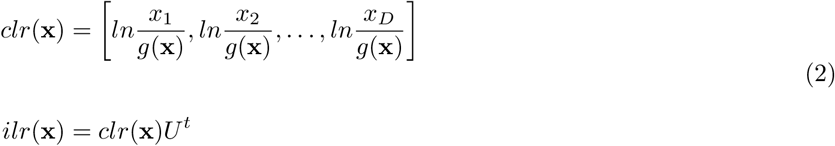

where 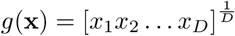 is the geometric mean of the composition and *U^t^* is a matrix which columns form an orthonormal basis of the centered log-ratio (clr) plane of x. Starting with a initial reference *r* (most proximal PAS, *r_0_* = x_1_), we iterate over each subsequent PAS (*t* = [x_2_,...,x*_D_*]) testing for DPU by fitting a linear model in a ilr-transformed sub-composition comprised of a reference and a target C[*r, t*], updating the reference to the target whenever the difference in compositions is not significant.

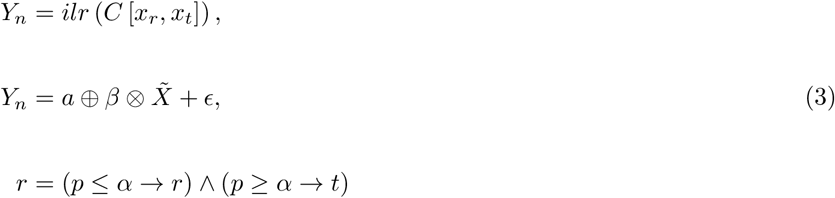

where C represents a closing operation in the sub-composition and *p* = *P(β* = 0) is the probability of the slope being 0.

## Simulation benchmark

To assess the performance of REPAC we used simFlux [25] to simulated fastq files of the dataset with 2 conditions and 5 replicates containing 5000 genes with two isoforms: a normal one (annotated) and one which 3’-UTR were extended by 1000 bp from the original gene model. Each replicates contained 5000 non-overlapping gene models that were randomly selected from the GENCODE 37 annotation and were composed of 2500 genes with longer/shorter isoforms equally distributed and 2500 genes without preferential isoform. The overall expression levels of the genes for each replicate were randomly assigned and the proportion of extended isoform usage was randomly assigned proportion *p* (0.1 ≥ *p* ≤ 0.35) in the first condition and 1 – p for the second condition. The fastq files were aligned with STAR with default parameters against the hg38 human genome. For the benchmark of REPAC, we quantified 50 bp windows located 100 bp upstream of the PAS (TTS of the isoforms) using featureCounts and the resulting matrix was used as input to REPAC to compare the two simulated conditions. Similarly, for DaPars[6] we generated wig files from the alignments and used them as input to DaPars along with a gtf file of the simulated isoforms. Finally, to benchmark QAPA[4] we quantified the 3’-UTR expression with salmon v1.6.0[9] using an index of 3’-UTR sequences extracted with the QAPA helper script and passed the quantification as input to QAPA. The absolute effect size of each method i.e., cFC, *ΔPDUI*, and ΔPAU for REPAC, Dapars, and QAPA, respectively was used as a score variable to compute the empirical ROC curves and AUC with the R/Bioconductor package ROCit.

## Resources and pre-processing

Both REPAC and QAPA rely on pre-annotated PAS to infer DPU. In this work, we used PAS annotations from the PolyAsite database for both mouse (mm10) and human (hg38) genomes. For each genome, we processed the PAS using *QAPA build* to incorporate PolyASite into the latest Ensembl annotations for each genome (v102 and v105 for mm10 and hg38, respectively) and select PAS overlapping with 3’-UTR regions. A total of 67509 and 85476 PAS coordinates were obtained and used in the downstream analysis for mice and humans, respectively. For the QAPA analysis, we extracted the 3’-UTR sequences using *QAPA* fasta. These sequences were used to build the index and perform the 3’-UTR expression quantification with salmon[9]. For the REPAC analysis, we generated a BED file with coordinates for 50bp windows located 50bp upstream of the PAS (herein referred to just as polyadenylation sites), which were used as input to the REPAC package. The whole transcriptome index for salmon[9] was built from the latest set of CDS and ncRNAs for the mouse genome from the Ensembl website to obtain the gene level quantification (see Technical implications of APA).

## Differential polyadenylation in activated B cells

Differential polyadenylation usage analyses were performed using REPAC and QAPA (v1.3)[4] to detect differential PAS usage between naive and activated B cells. The REPAC analysis was carried out using the REPAC package to query recount3 tracks and quantify PAS to obtain an expression matrix. Next, we filtered low expressed sites (*counts <* 10) and low expressed genes (*counts <* 30) from the analysis using the salmon whole transcriptome quantification (see Technical implications of APA). Finally, we estimated the cFC between PAS by fitting a linear model on the ilr-transformed compositions. Compositions with *|cFC*| ≥ 0.25 and adjusted p-value ≤ 0.05 were considered shortening or lengthening events if their cFC were negative or positive, respectively. For the GSEA analysis, we ranked the results of REPAC by t-statistics and tested the MSigDB GO Biological process collection for negative enrichment using a Monte Carlo adaptive multilevel splitting approach, implemented in the fgsea package[26]. The results of GSEA were collapsed with fgsea collapsePathways function to reduce redundancy.

For the QAPA analysis, we used the QAPA quant option to load 3’-UTR expression estimates from salmon and compute the PAU’s for each sample. Low expressed 3’-UTRs were filtered (TPM ≤ 5) and the *ΔPPAU* was computed as the difference between the average PPAU for each condition. Finally, genes with *|ΔPPAU*| ≥ 20 were considered shortening or lengthening events if their *ΔPPAU* were negative or positive, respectively. Enrichment analysis of GO BP was conducted in the same manner as REPAC, but ranking the genes by *ΔPPAU* instead.

## Technical implications of APA in downstream analyses

To assess the impact 3’-US events can have in gene expression estimations we obtained the raw data from SRA (SRP048707 [12]) and estimated the transcripts expression levels with salmon[9]. The transcripts estimates were summarized at gene level with tximport[24] setting the argument “countsFromAbundance” to “no” (raw expression values) and “lengthScaledTPM” (isoform length-corrected values). For each estimate, low count genes (< 5 counts) were filtered and the remaining genes were normalized with the trimmed mean of the M-Values method. A generalized linear model approach coupled with empirical Bayes moderation of standard errors and voom precision weights[27, 28] was used to detect deferentially expressed genes between the selected contrasts. Adjusted p-values controlling for multiple hypothesis testing were performed using the Benjamini-Hochberg method[29]. Next, the ratio between the fold-change of the two results for the genes with significant DPU was used to estimate the impact on downstream analysis.

## Analysis of GTEx tissues

To evaluate how well REPAC can scale with large data sets we randomly selected a subset of 20, 100, 200, and 400 brains (cortex) and testis samples from the GTEx tracks of recount3[8]. For each subset, we quantified the PAS using the REPAC function *create_pa_rse* and recorded the time taken to quantify each subset. Next, We filtered PASs that were lowly expressed (counts ≤ 10) and estimated the cFC between PAS by fitting a linear model on the ilr-transformed compositions. PAS with a *|cFC*| ≥ 0.25 and *FDR ≤* 0.05 were considered shortening or lengthening events if their cFC were negative or positive, respectively. GSEA analysis was carried out by ranking the results of REPAC by t-statistics and testing the MSigDB hallmarks collection with the fgsea package[26].

## Declarations

Ethics approval and consent to participate

Not applicable.

## Consent for publication

Not applicable.

## Availability of data and materials

All raw data used in this work are available at the Sequence Read Archive (SRA) (https://www.ncbi.nlm.nih.gov/sra) study SRP048707. Counts estimates from GTEx are available through the recount3 and REPAC R packages. The REPAC R package can be obtained at https://github.com/eddieimada/REPAC. All scripts to reproduce the analyses in this manuscript, including simulation, are available at https://github.com/eddieimada/REPAC_paper.

## Author’s contributions

LM and ELI conceived the idea, design the study and interpreted the results; ELI performed the analysis; CW and BL provided data and tools; All authors wrote, reviewed and approved the manuscript.

## Acknowledgements

recount3 is hosted on SciServer, a collaborative research environment for large-scale data-driven science. It is being developed at, and administered by, the Institute for Data Intensive Engineering and Science (IDIES) at Johns Hopkins University. SciServer is funded by the National Science Foundation Award ACI-1261715. For more information about SciServer, visit http://www.sciserver.org/.

## Figures

### Additional Files

Additional file 1 – Top 10 DPU genes detected by QAPA

3’-UTR coverage across Naive and LPS-activated B cells for the top 10 APA events detected by QAPA.

Additional file 2 – Top 10 DPU genes detected by REPAC

3’-UTR coverage across Naive and LPS-activated B cells for the top 10 APA events detected by REPAC.

Additional file 3 – REPAC differential polyadenylation usage results.

Additional file 4 – GSEA tables for REPAC and QAPA analysis

