## Supplementary figures and images for "Unleashing alternative polyadenylation analyses with REPAC" (from another version of this preprint: version 1, 2022-03-16)

### Additional File 1

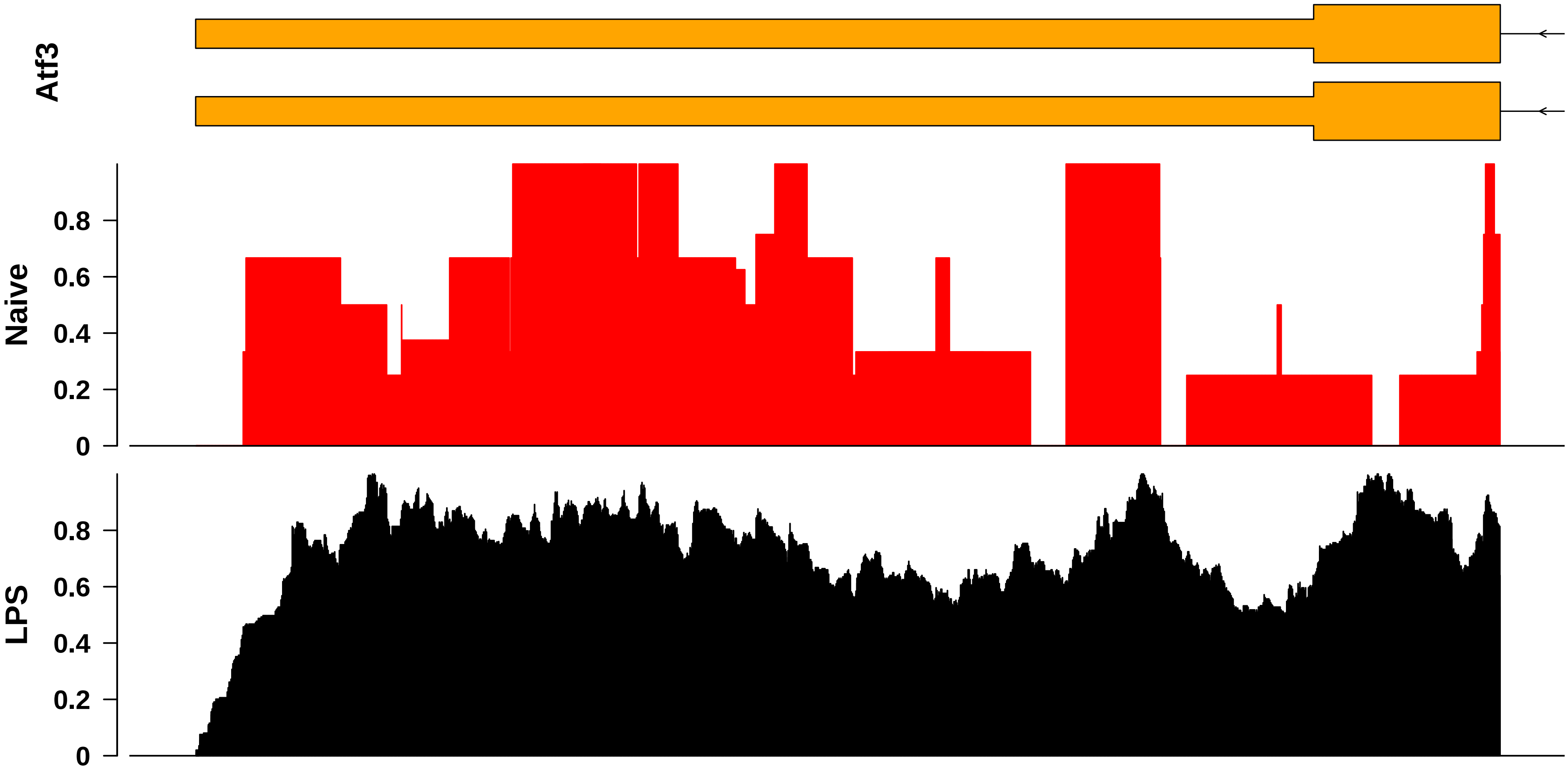

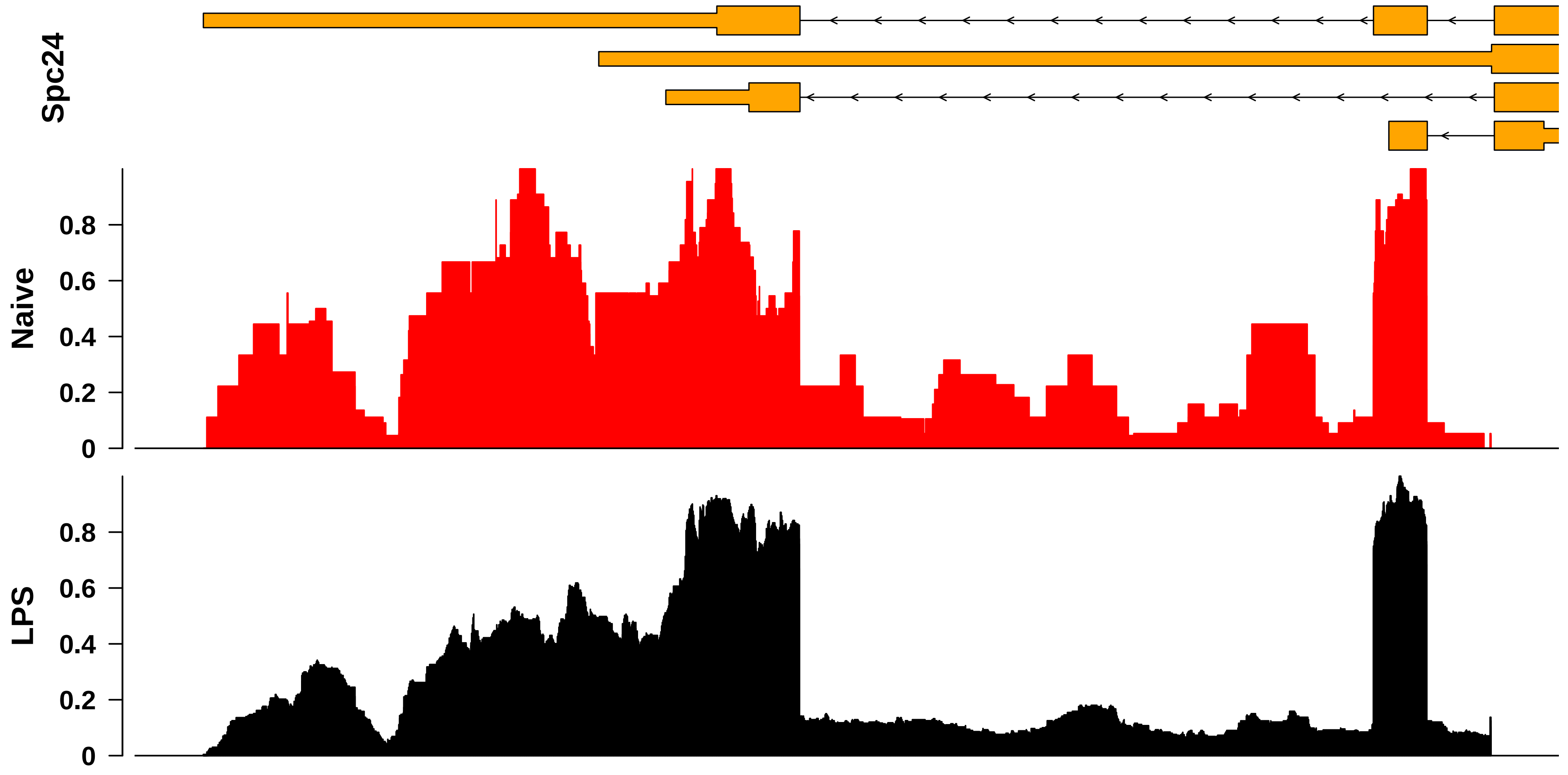

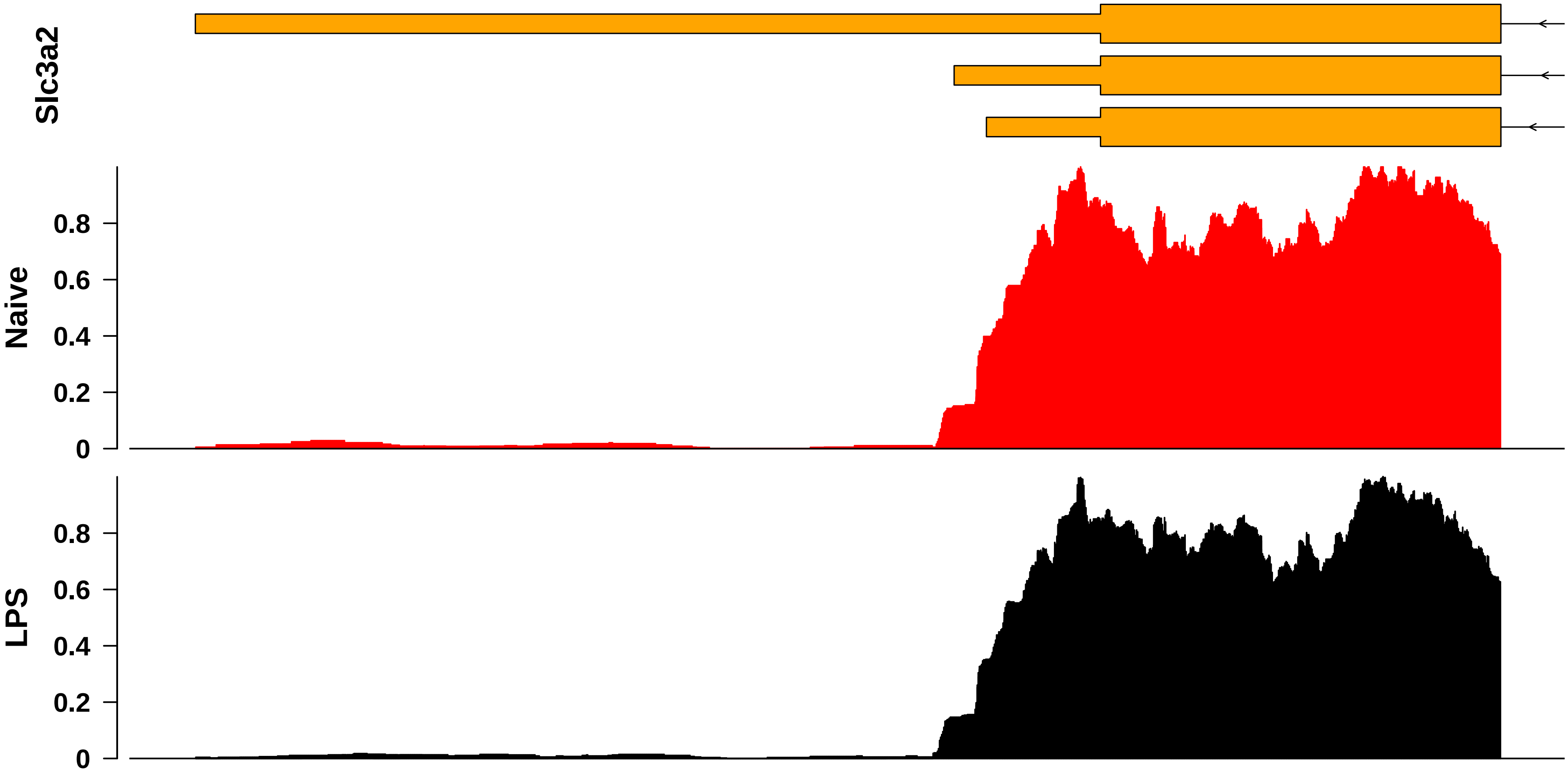

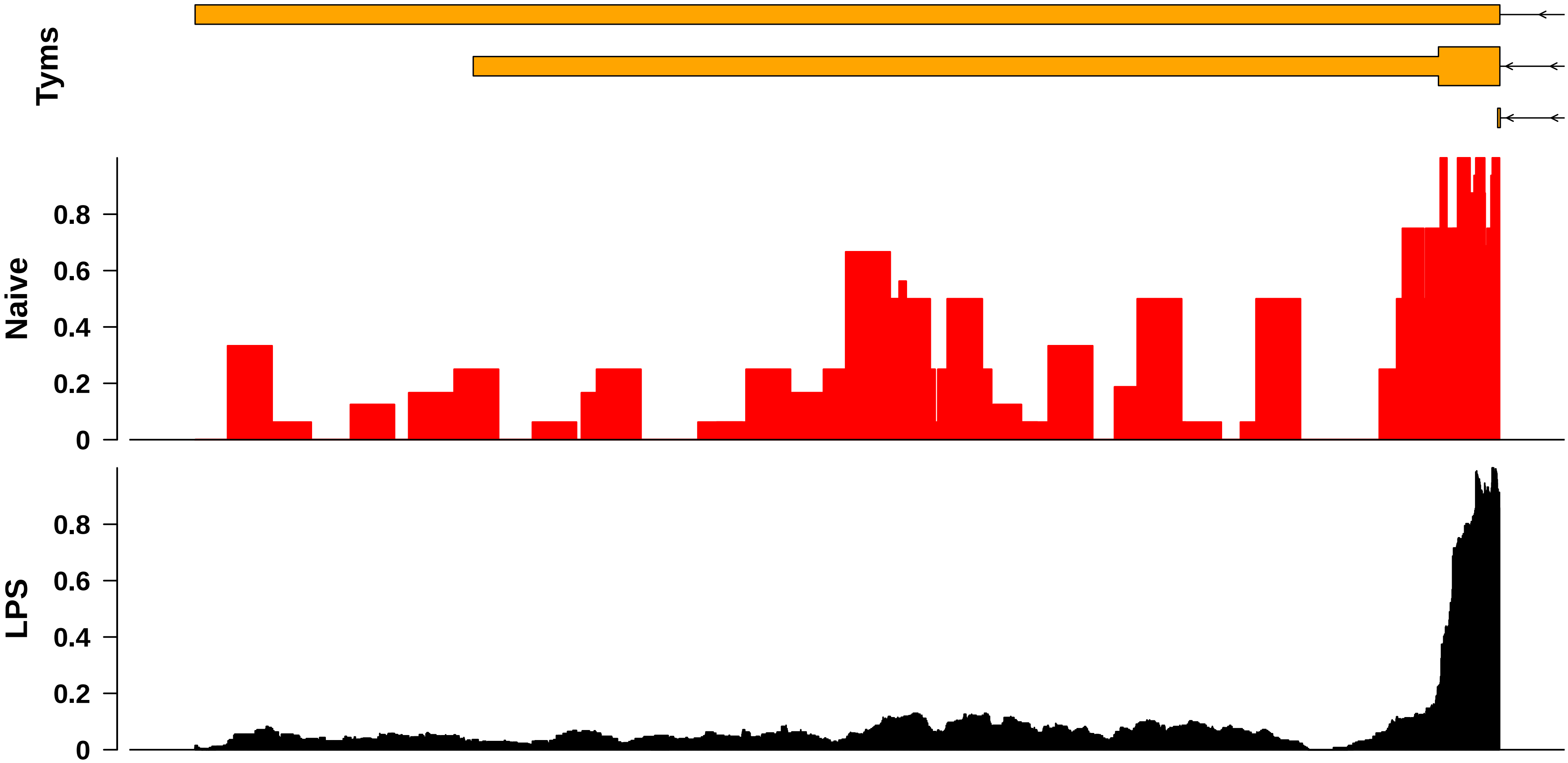

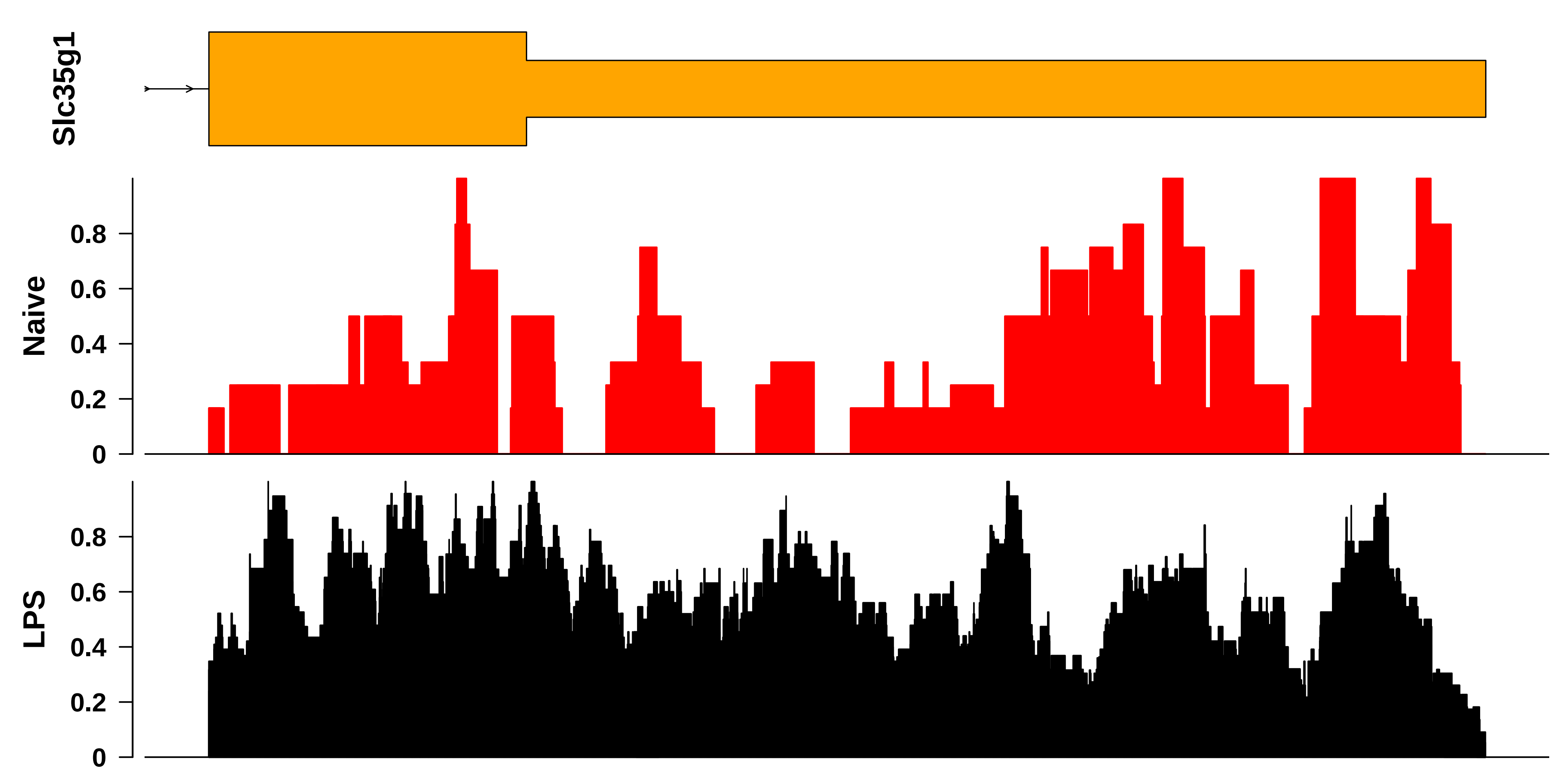

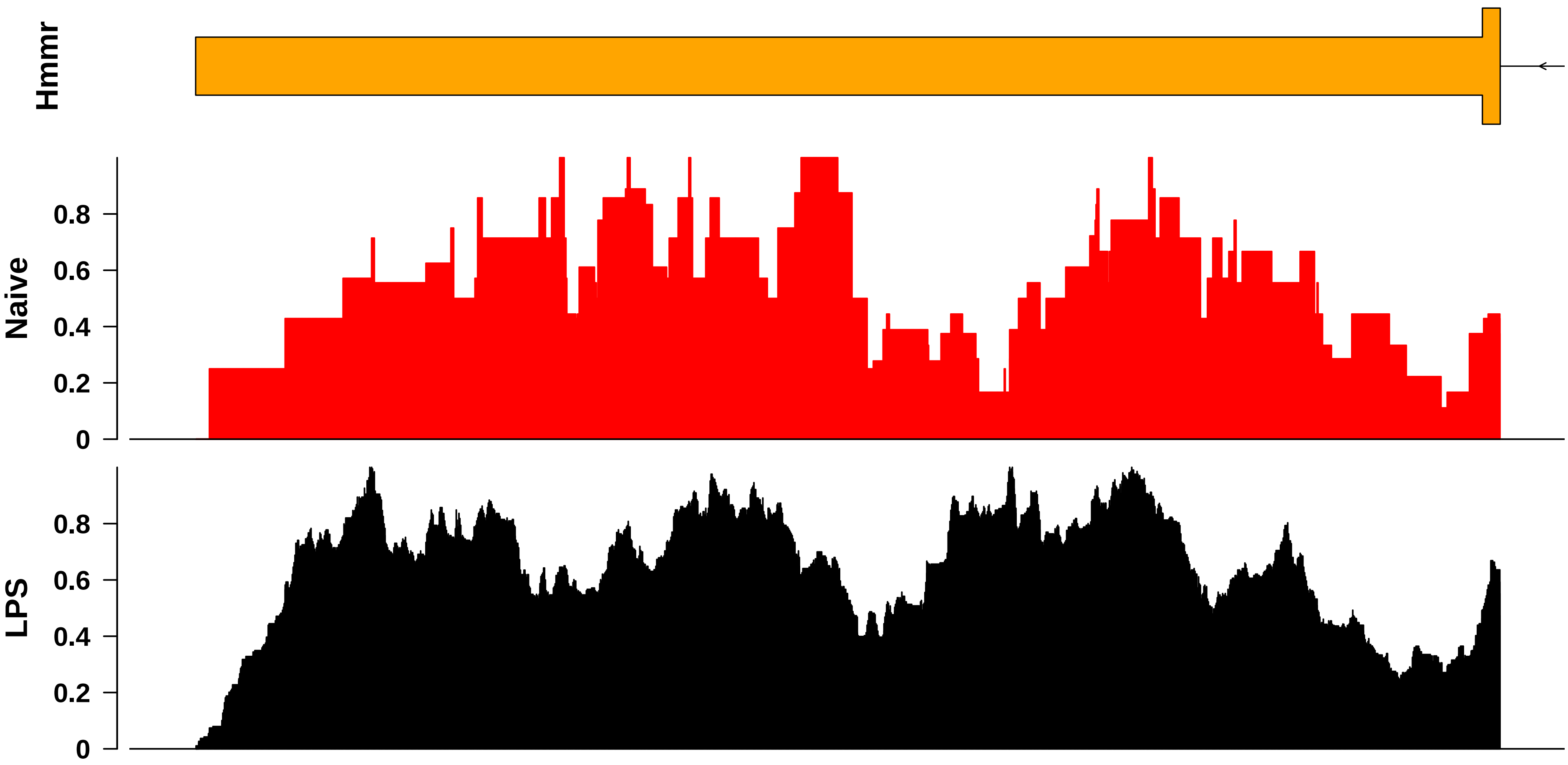

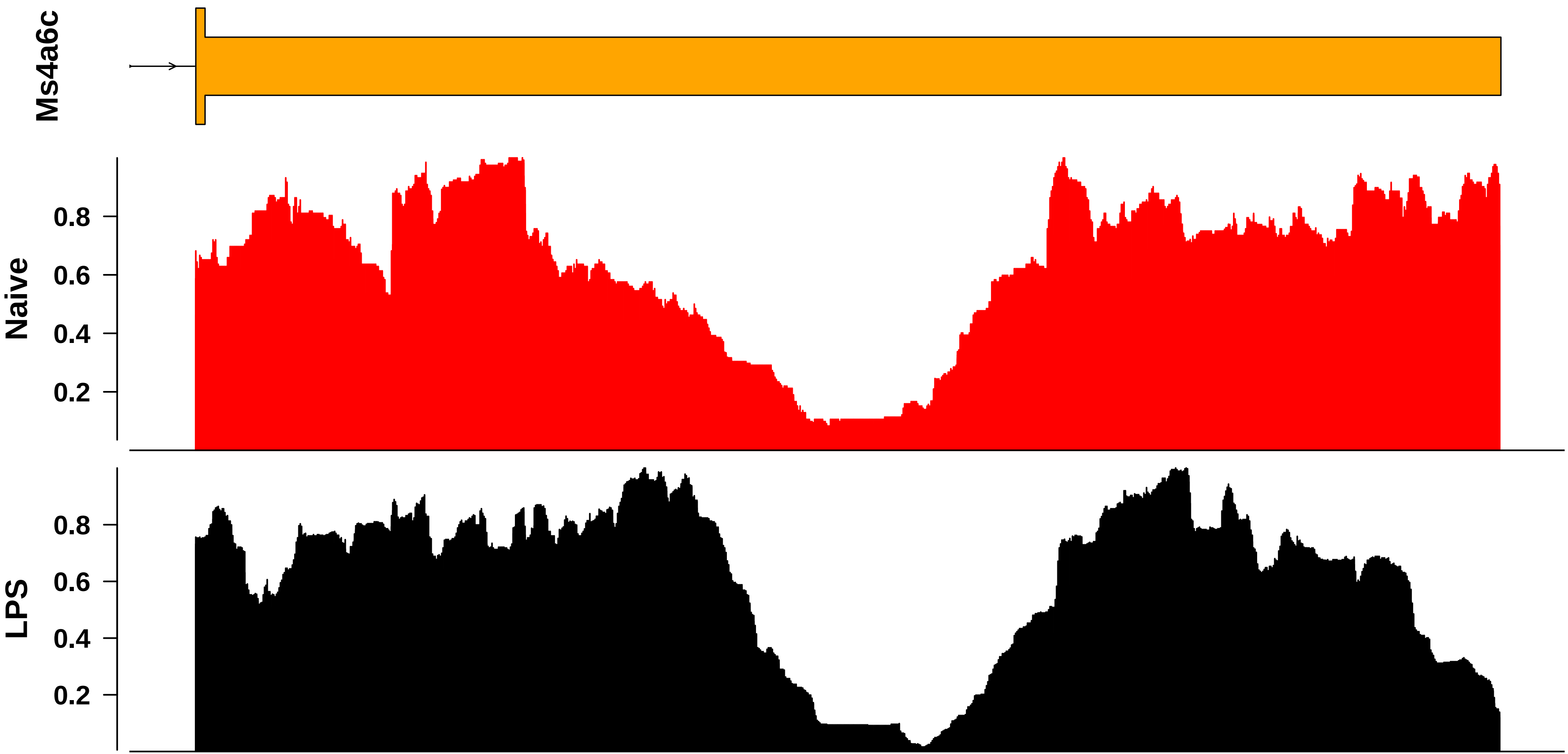

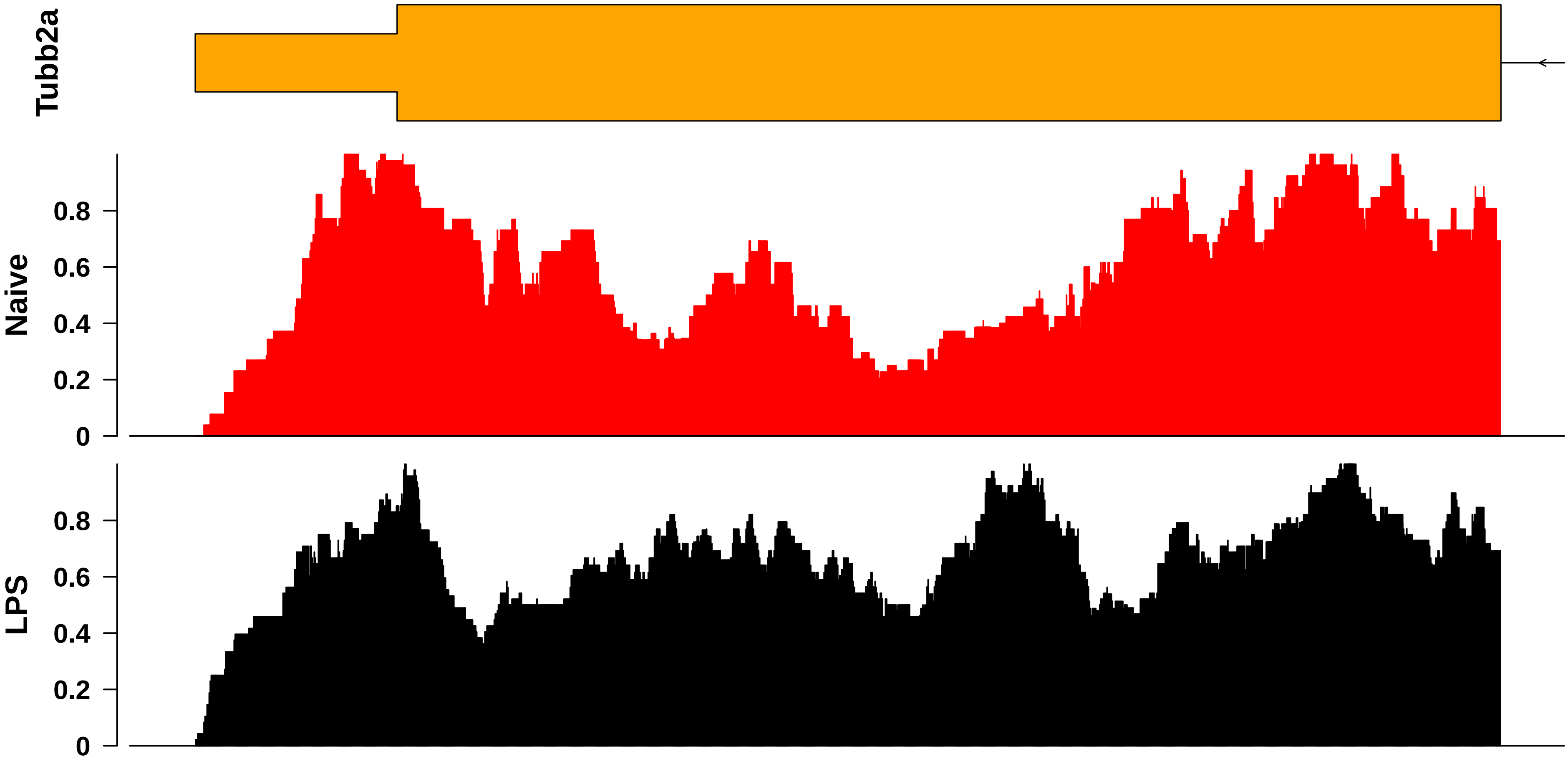

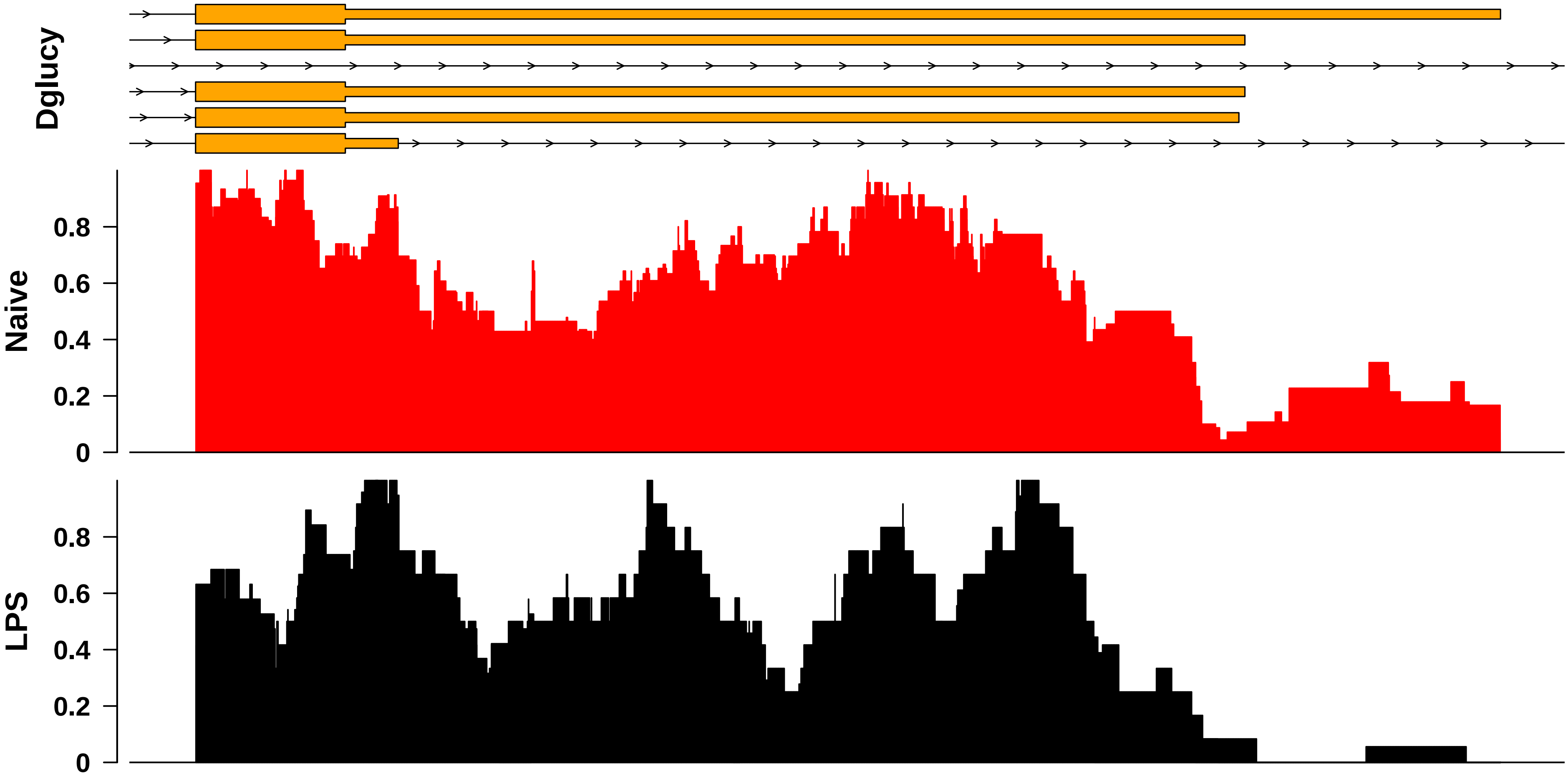

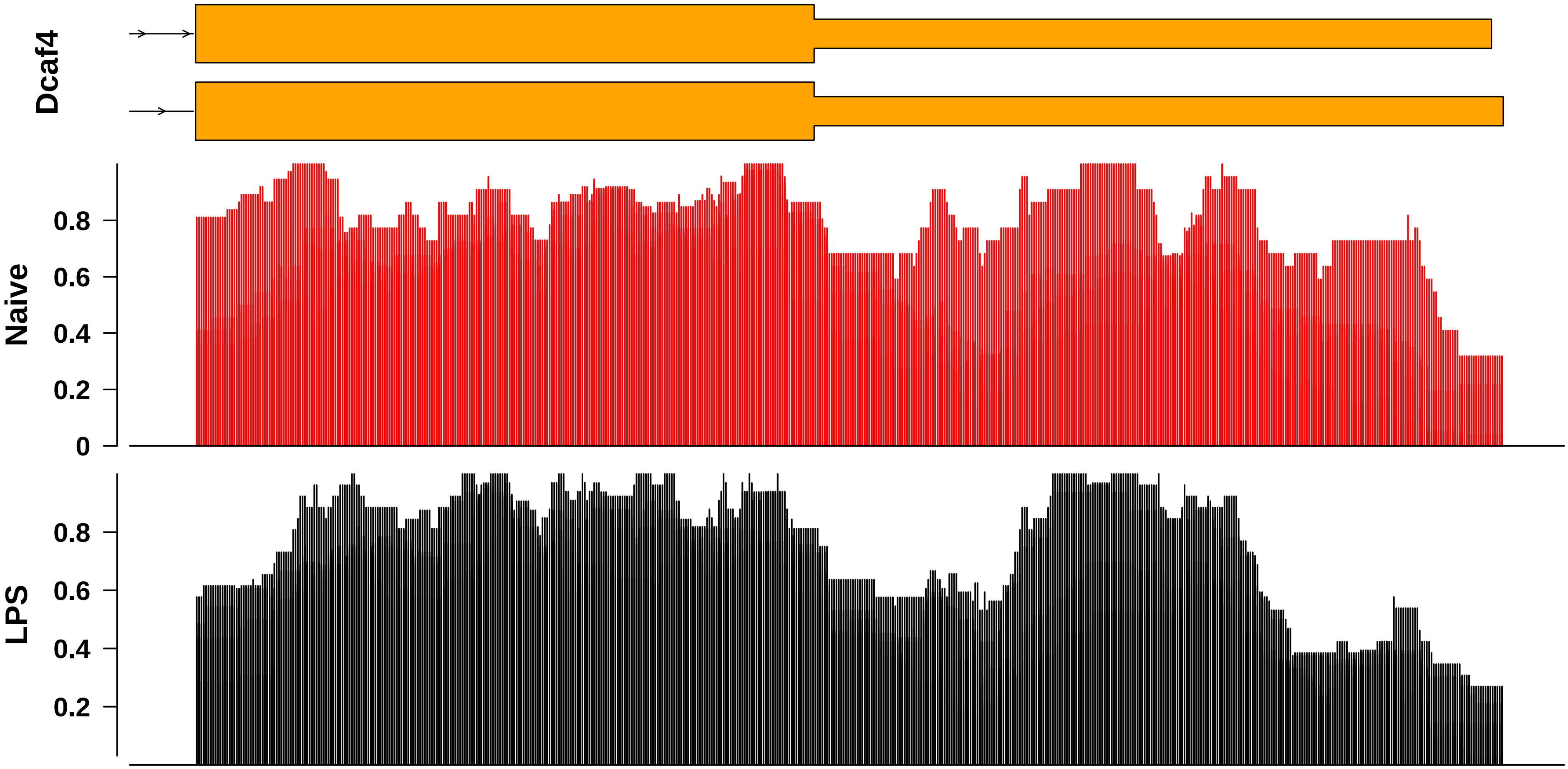

### Additional File 2

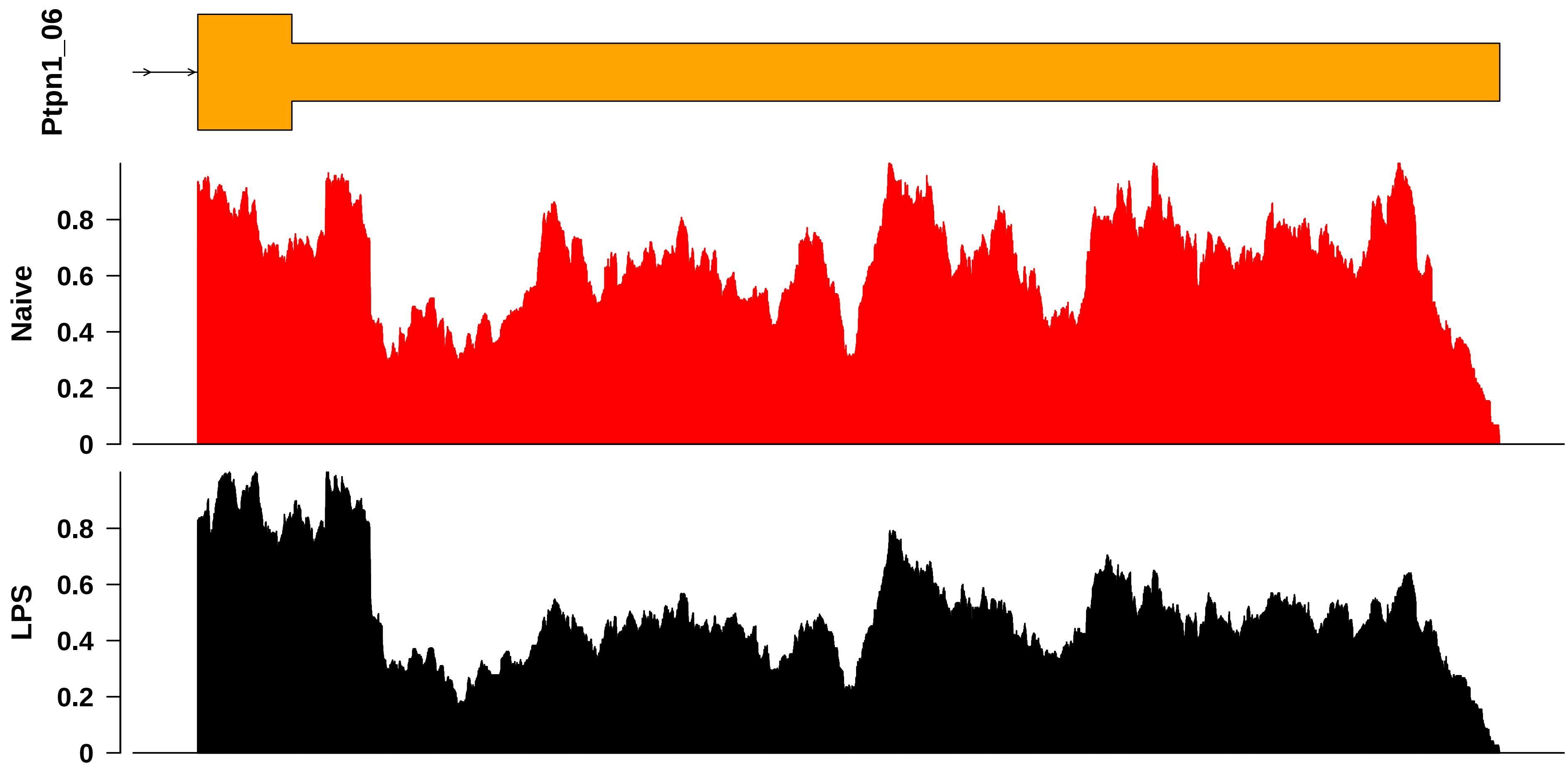

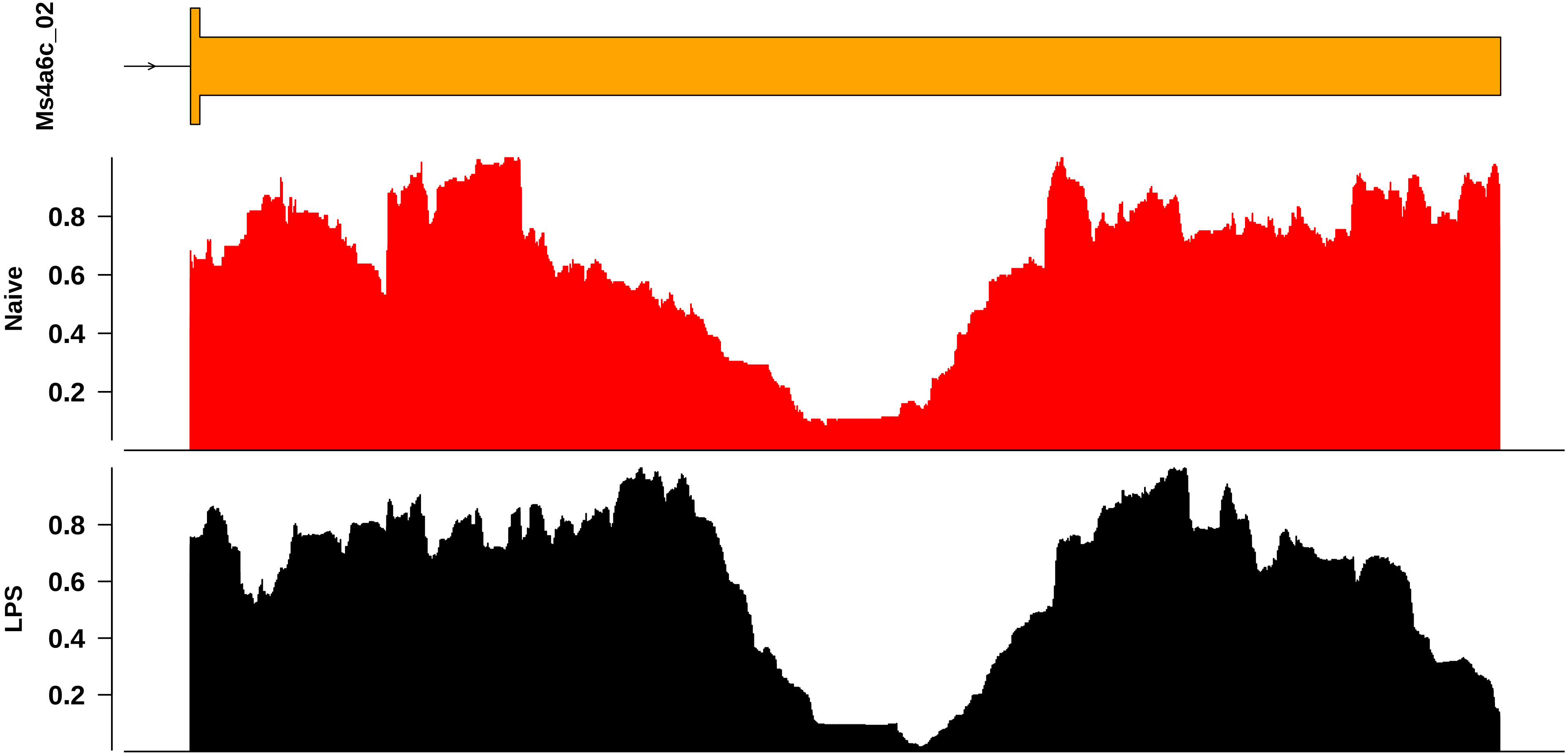

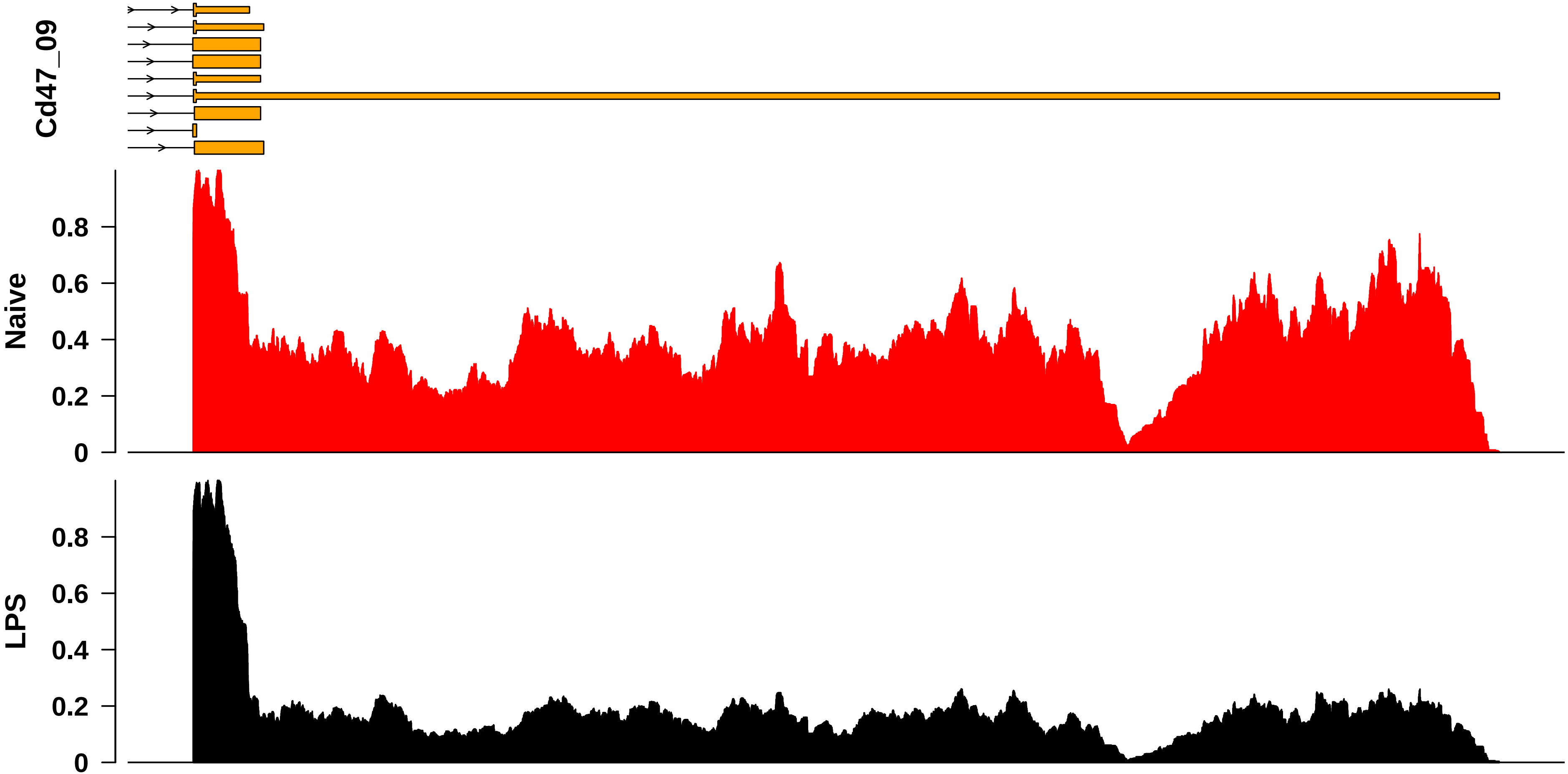

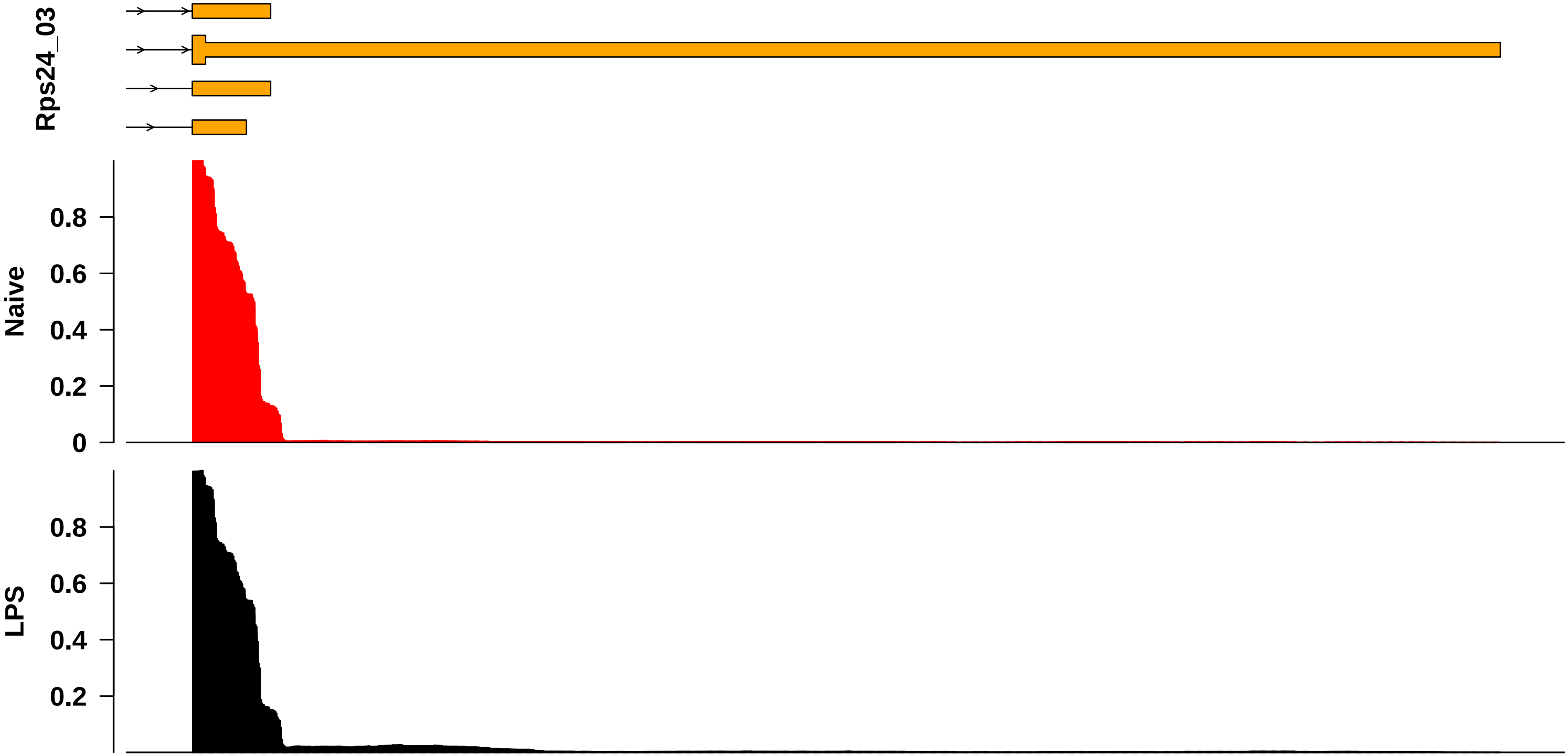

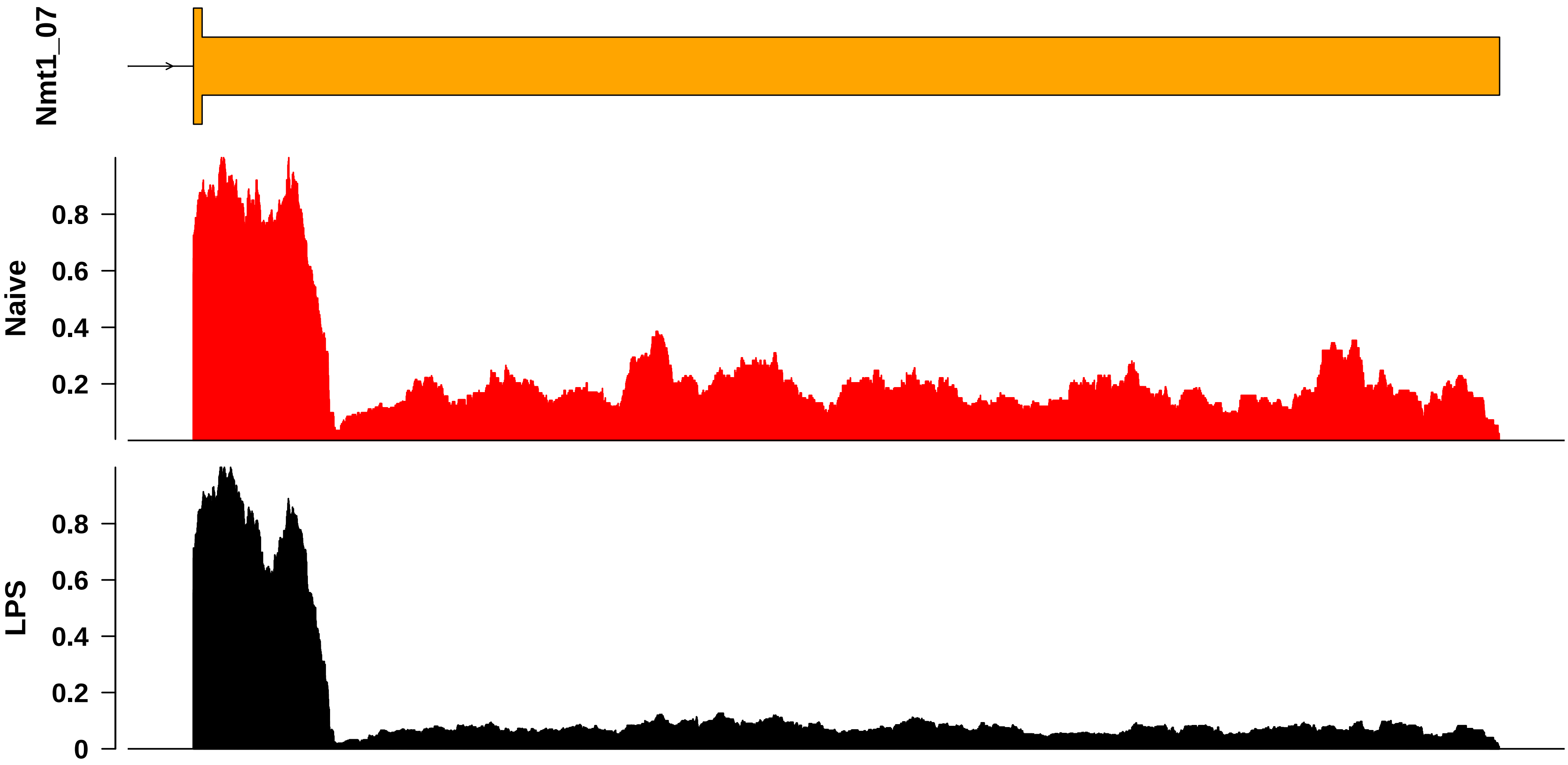

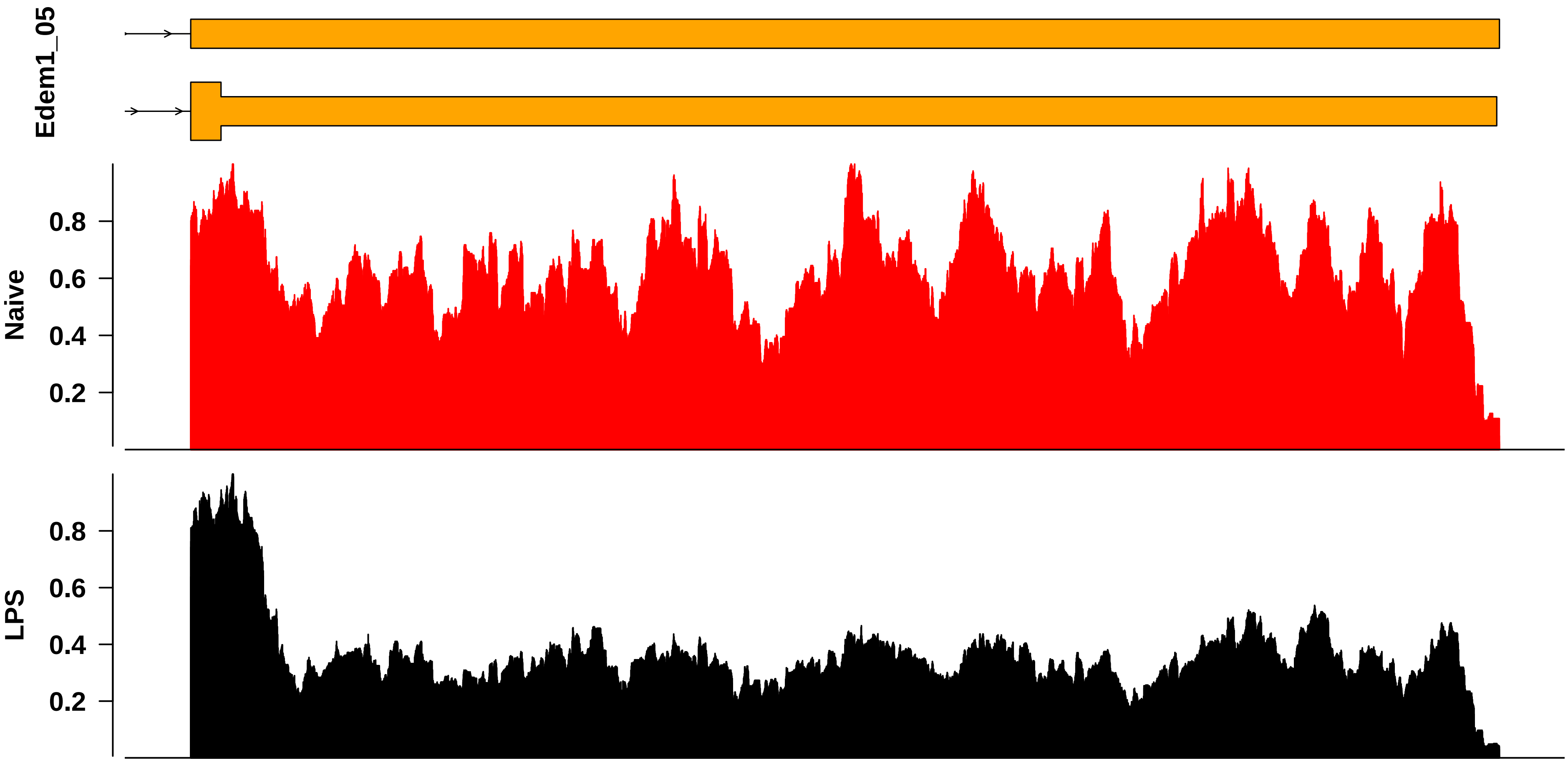

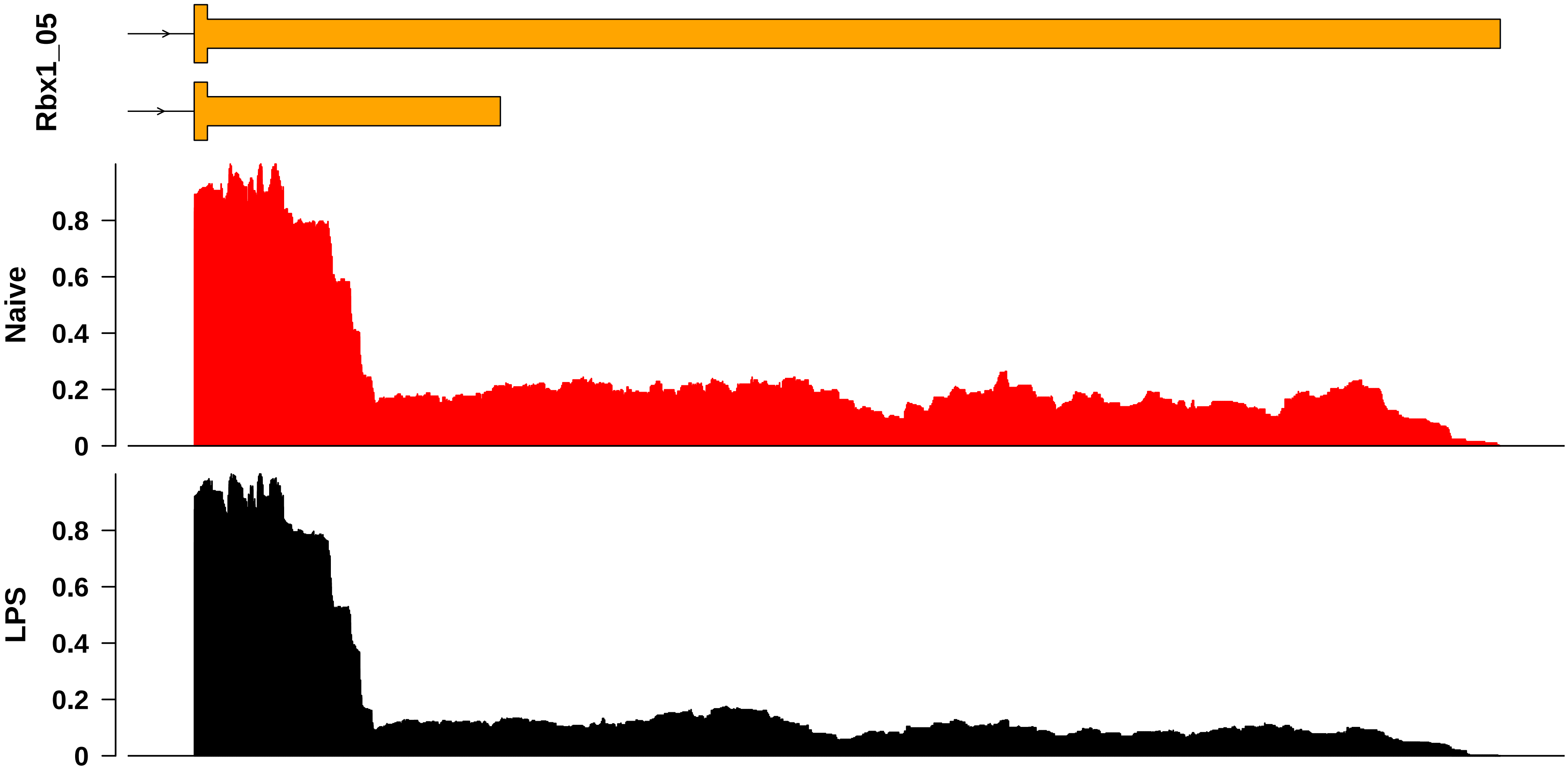

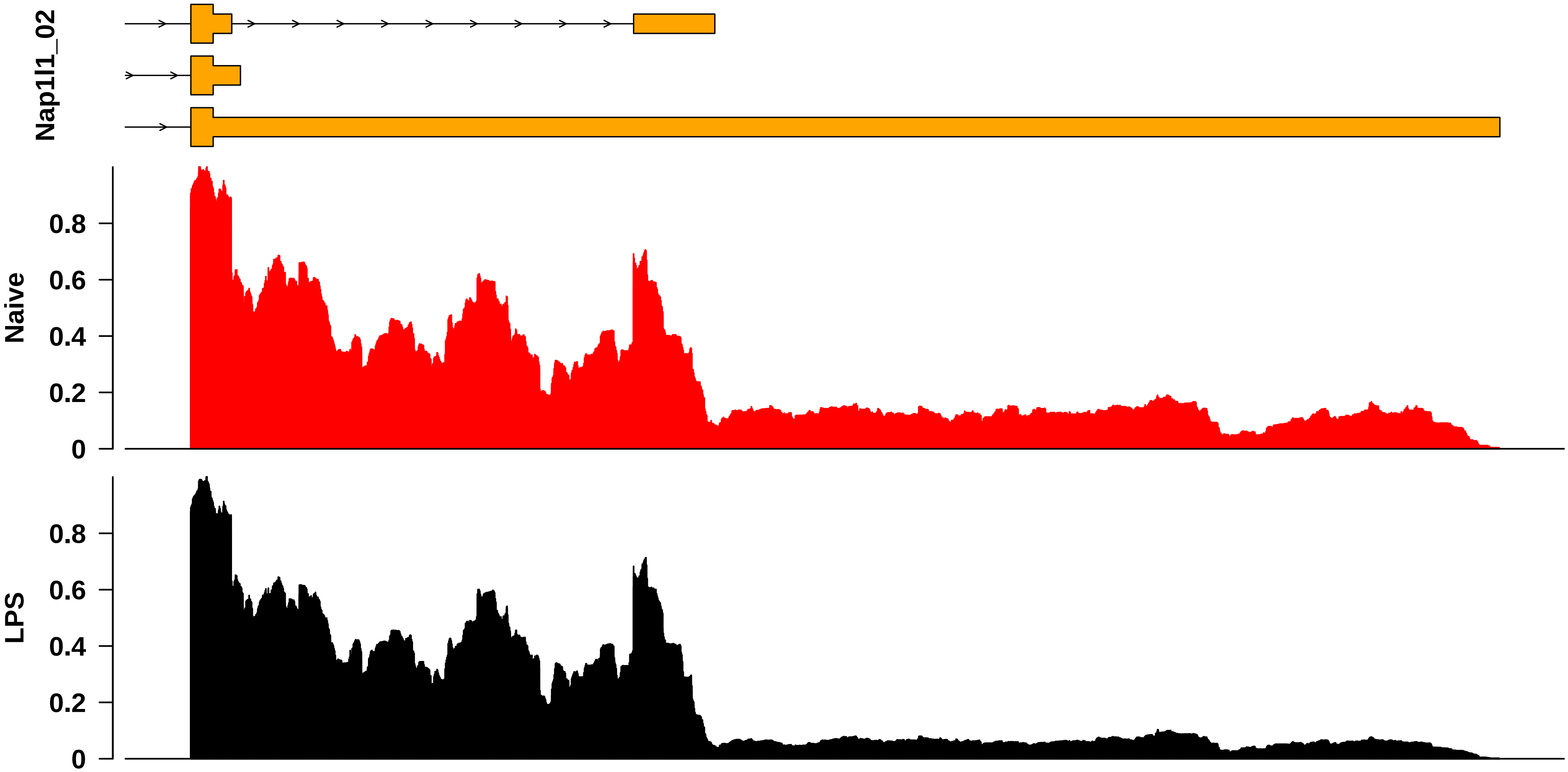

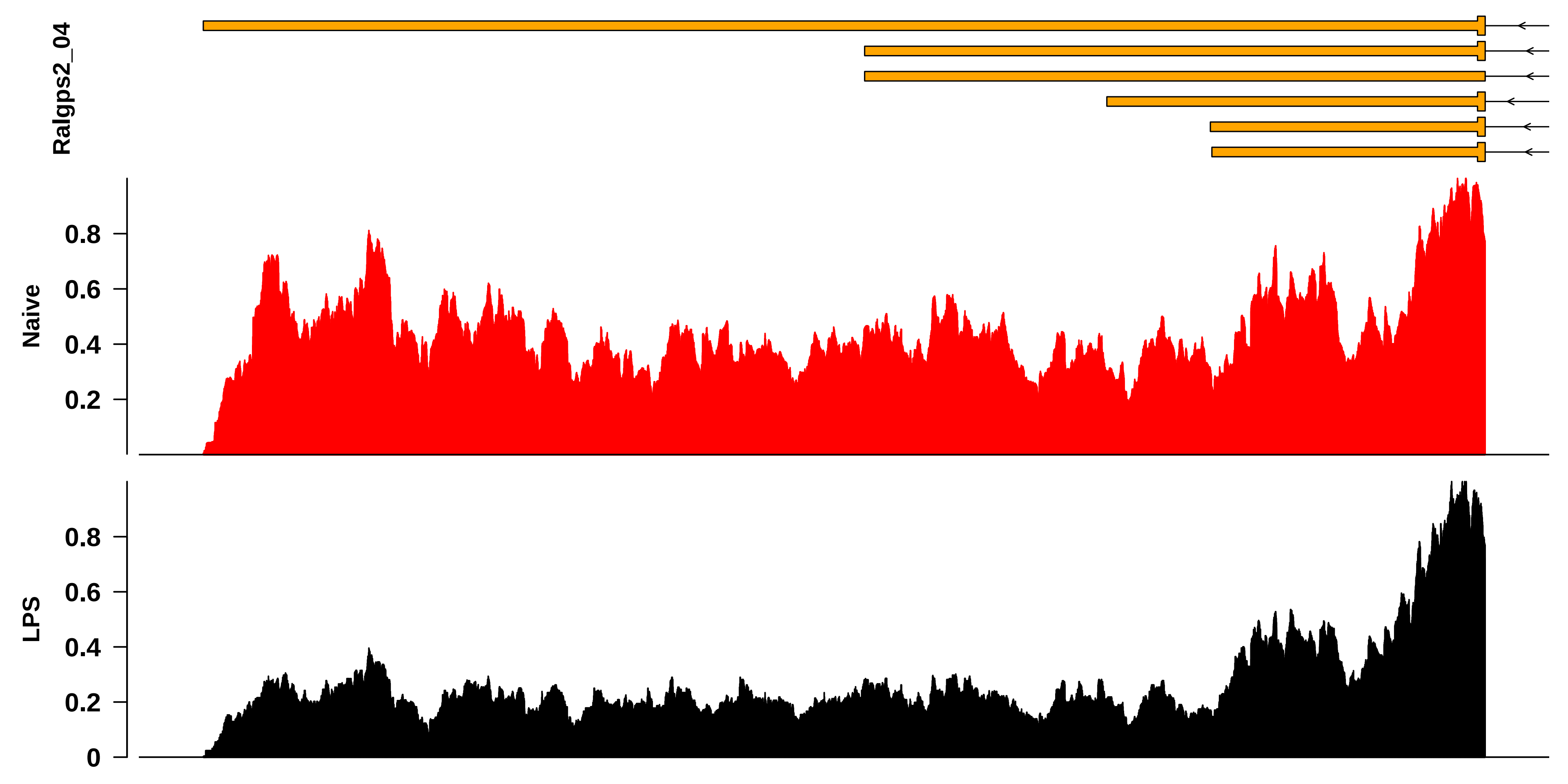

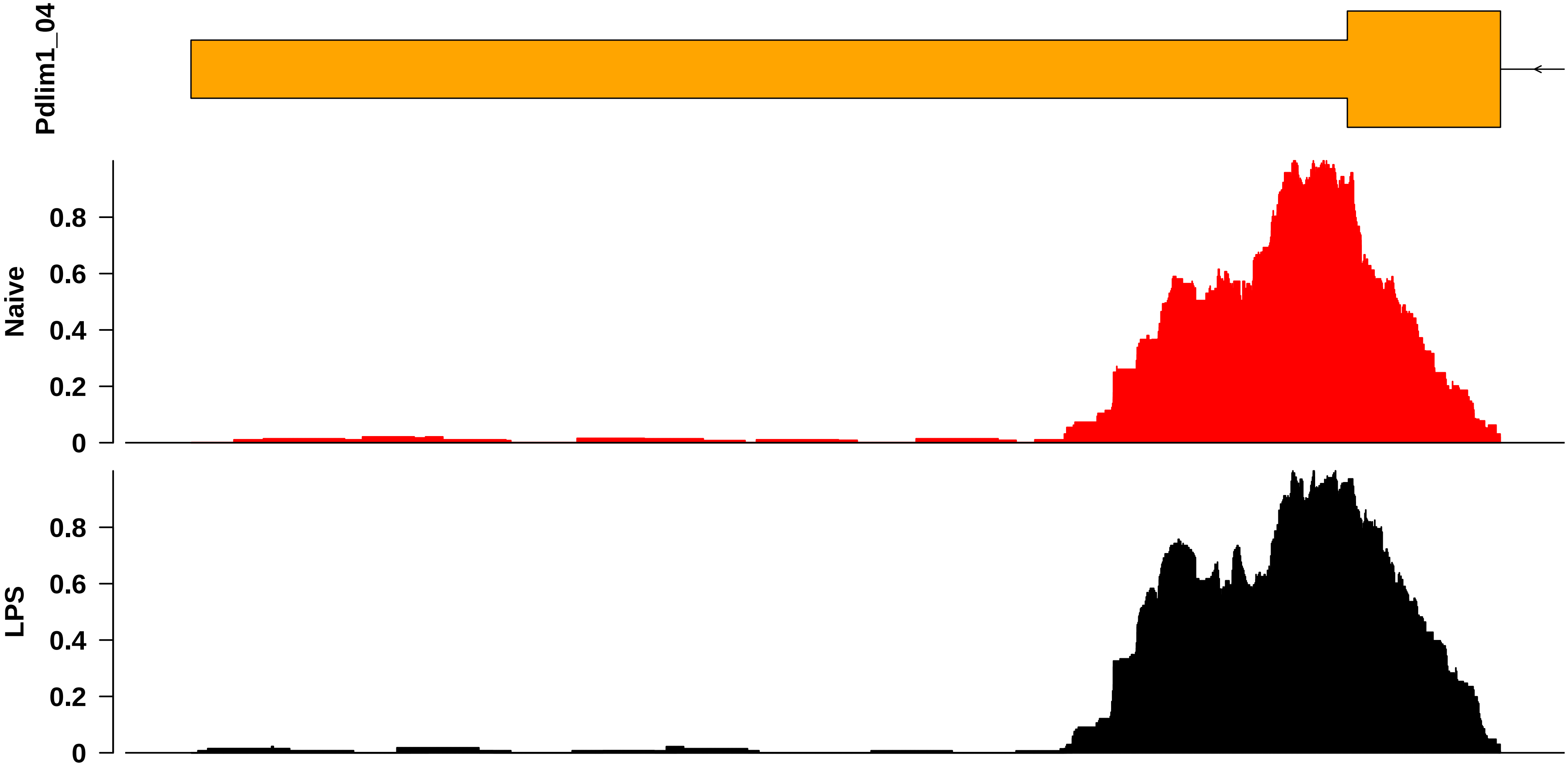
